# Conserved Neutralizing Epitopes on the N-Terminal Domain of Variant SARS-CoV-2 Spike Proteins

**DOI:** 10.1101/2022.02.01.478695

**Authors:** Zijun Wang, Frauke Muecksch, Alice Cho, Christian Gaebler, Hans-Heinrich Hoffmann, Victor Ramos, Shuai Zong, Melissa Cipolla, Briana Johnson, Fabian Schmidt, Justin DaSilva, Eva Bednarski, Tarek Ben Tanfous, Raphael Raspe, Kaihui Yao, Yu E. Lee, Teresia Chen, Martina Turroja, Katrina G. Milard, Juan Dizon, Anna Kaczynska, Anna Gazumyan, Thiago Y. Oliveira, Charles M. Rice, Marina Caskey, Paul D. Bieniasz, Theodora Hatziioannou, Christopher O. Barnes, Michel C. Nussenzweig

**Affiliations:** Laboratory of Molecular Immunology, The Rockefeller University, New York, NY 10065, USA; Laboratory of Retrovirology, The Rockefeller University, New York, NY 10065, USA; Laboratory of Virology and Infectious Disease, The Rockefeller University, New York, NY, USA; Department of Biology, Stanford University, Stanford, CA 94305, USA; Howard Hughes Medical Institute; Chan Zuckerberg Biohub, San Francisco, CA 94158, USA

## Abstract

SARS-CoV-2 infection or vaccination produces neutralizing antibody responses that contribute to better clinical outcomes. The receptor binding domain (RBD) and the N-terminal domain (NTD) of the spike trimer (S) constitute the two major neutralizing targets for the antibody system. Neutralizing antibodies targeting the RBD bind to several different sites on this domain. In contrast, most neutralizing antibodies to NTD characterized to date bind to a single supersite, however these antibodies were obtained by methods that were not NTD specific. Here we use NTD specific probes to focus on anti-NTD memory B cells in a cohort of pre-omicron infected individuals some of which were also vaccinated. Of 275 NTD binding antibodies tested 103 neutralized at least one of three tested strains: Wuhan-Hu-1, Gamma, or PMS20, a synthetic variant which is extensively mutated in the NTD supersite. Among the 43 neutralizing antibodies that were further characterized, we found 6 complementation groups based on competition binding experiments. 58% targeted epitopes outside the NTD supersite, and 58% neutralized either Gamma or Omicron, but only 14% were broad neutralizers. Three of the broad neutralizers were characterized structurally. C1520 and C1791 recognize epitopes on opposite faces of the NTD with a distinct binding pose relative to previously described antibodies allowing for greater potency and cross-reactivity with 7 different variants including Beta, Delta, Gamma and Omicron. Antibody C1717 represents a previously uncharacterized class of NTD-directed antibodies that recognizes the viral membrane proximal side of the NTD and SD2 domain, leading to cross-neutralization of Beta, Gamma and Omicron. We conclude SARS-CoV-2 infection and/or Wuhan-Hu-1 mRNA vaccination produces a diverse collection of memory B cells that produce anti-NTD antibodies some of which can neutralize variants of concern. Rapid recruitment of these cells into the antibody secreting plasma cell compartment upon re-infection likely contributes to the relatively benign course of subsequent infections with SARS-CoV-2 variants including omicron.

## Introduction

Several independent studies purified anti-SARS-CoV-2 specific B cells from infected or vaccinated individuals using soluble spike (S) protein as a bait. In all cases the neutralizing antibodies obtained by this method targeted the RBD most frequently and were generally more potent than those targeting the NTD (Kreer et al., 2020; Liu et al., 2020; Zost et al., 2020b).

Further characterization of the neutralizing antibodies to RBD showed that infected or vaccinated humans produce a convergent set of neutralizing antibodies dominated by specific Ig heavy chain variable (IGVH) regions (Barnes et al., 2020b; Brouwer et al., 2020; Robbiani et al., 2020; Wang et al., 2021d; Yuan et al., 2020). Structural analysis of the interaction between these antibodies and the RBD revealed four classes of anti-RBD antibodies each of which can interfere with the interaction between this domain and ACE2 the cellular receptor for the SARS-CoV-2 S protein (Barnes et al., 2020a; Yuan et al., 2020). The most frequently occurring anti-RBD antibodies select for resistance mutations in *in vitro* experiments that are also found in variants of concern (Baum et al., 2020; Greaney et al., 2021; Wang et al., 2021d; Weisblum et al., 2020). These include the K417N, E484A and N501Y changes that are found in Omicron, the most recently emerging variant of concern (Callaway, 2021). However, anti-RBD antibodies that remain resistant to these changes evolve over time in the memory B cell compartment of both naturally infected and vaccinated individuals over time (Cho et al., 2021; Muecksch et al., 2021; Wang et al., 2021c).

Less is known about anti-NTD antibody responses. In contrast to RBD, the anti-NTD neutralizing antibodies obtained to date primarily target a single supersite consisting of variable loops flanked by glycans (Amanat et al., 2021; Cerutti et al., 2021b; Chi et al., 2020; Dussupt et al., 2021; Haslwanter et al., 2021; Li et al., 2021; Liu et al., 2021; McCallum et al., 2021a; Planas et al., 2021b; Suryadevara et al., 2021b; Voss et al., 2021). Residues in the supersite are mutated in several variants of concern including Beta, Gamma and Omicron, the latter of which carries 3 deletions, 4 amino acid substitutions and an insertion in the NTD that render antibodies to the supersite infective (Callaway, 2021; Faria et al., 2021; Tegally et al., 2021). Residues in this site are also mutated in the S protein of PMS20, a synthetic construct that is highly antibody resistant and chimeric proteins build from WT and PMS20 proteins show that NTD-specific antibodies are an important component of the neutralizing activity in convalescent and vaccine recipient plasma (Schmidt et al., 2021c). NTD supersite mutations are therefore likely to contribute to the poor plasma neutralizing activity against the Omicron variant in individuals that received 2 doses of currently available vaccines or convalescent individuals exposed to pre-Omicron variants of SARS-CoV-2. Nevertheless, boosting with currently available mRNA vaccines elicits high levels of plasma Omicron neutralizing antibodies(Dejnirattisai et al., 2022; Gruell et al., 2022; Rossler et al., 2022; Schmidt et al., 2021b; Wu et al., 2022), and vaccinated individuals are protected from serious disease following Omicron infection. Whether additional anti-NTD omicron neutralizing epitopes exist, and how memory B cell anti-NTD neutralizing responses evolve over time is poorly understood.

To focus on the development and evolution of the human antibody response to NTD we studied a cohort of SARS-CoV-2 convalescent and/or mRNA vaccinated individuals using the isolated NTD domain as a probe to capture memory B cells producing antibodies specific to this domain.

## Results

To examine the development of anti-NTD antibodies we studied a previously described longitudinal cohort of individuals sampled 1.3 and 12 months after infection, some of whom received an mRNA vaccine approximately 40 days before the 12-month study visit ((Wang et al., 2021c) and Table S1). All individuals were infected between 1 April and 8 May of 2020, before the emergence of Omicron. Antibody reactivity in plasma to the isolated Wuhan-Hu-1 and Omicron NTD was measured by ELISA (Figure S1). There was no association between IgM, IgG or IgA anti-NTD Wuhan-Hu-1 or Omicron ELISA titers and age, sex, symptom severity, duration of symptoms, persistence of symptoms, or time between symptom onset and the first clinic visit (Figure S2).

In convalescent individuals that had not been vaccinated anti-Wuhan-Hu-1 NTD IgG reactivity was not significantly different between the 1.3- and 12-month time points (Figure S1A). In contrast, IgG reactivity to the Omicron NTD dropped significantly between the 2 time points (Figure S1B). Notably, there was no correlation between NTD and RBD IgG ELISA reactivity or NTD IgG ELISA reactivity and plasma geometric mean half-maximal neutralizing titers (NT_50_s, see below) in convalescent individuals that had not been vaccinated (Figure S3). Vaccination of convalescent individuals resulted in significantly increased IgG ELISA reactivity to both Wuhan-Hu-1 and Omicron NTDs with a positive correlation between NTD and RBD ELISA, and NTD and plasma NT_50_ (Figures S3A and S3B, (Schmidt et al., 2021a)). IgM and IgA anti-NTD reactivity in plasma was relatively low in all individuals tested, and only IgA was boosted with vaccination (Figures S1C and S1D). We conclude that anti-NTD ELISA reactivity to both Wuhan-Hu-1 and Omicron is enhanced by vaccination in convalescent individuals.

### Memory anti-NTD Antibodies

Nearly all anti-NTD antibodies characterized to date were obtained using the intact S protein to capture specific memory B cells. Neutralizing anti-NTD antibodies obtained by this method represent a small subset of the total anti-S antibodies and in most cases, they target a single carbohydrate flanked epitope that faces away from the cell membrane and is mutated in several variants of concern including Omicron (Amanat et al., 2021; Cerutti et al., 2021b; Chi et al., 2020; Dussupt et al., 2021; Haslwanter et al., 2021; Li et al., 2021; Liu et al., 2021; McCallum et al., 2021a; Planas et al., 2021b; Suryadevara et al., 2021b; Voss et al., 2021). To focus on NTD we used a combination of soluble Wuhan-Hu-1 and Gamma NTD proteins to identify memory B cells producing antibodies specific for this domain.

Convalescent individuals that had not been vaccinated showed a small but significant decrease in the number of anti-NTD memory B cells between the 1.3- and 12-month time points, however, it was boosted by vaccination (Figures 1A and S4). We obtained 914 anti-NTD antibody sequences from 3 non-vaccinated and 3 vaccinated convalescent individuals assayed at the 2 time points (Figure 1B and Table S2). Like the anti-RBD response, different individuals made closely related antibodies to NTD among which antibodies encoded by VH1-24, VH3-30, VH3-33, VH4-4, and VH4-39 were over-represented (Figures 1C and S5).

**Figure 1.**
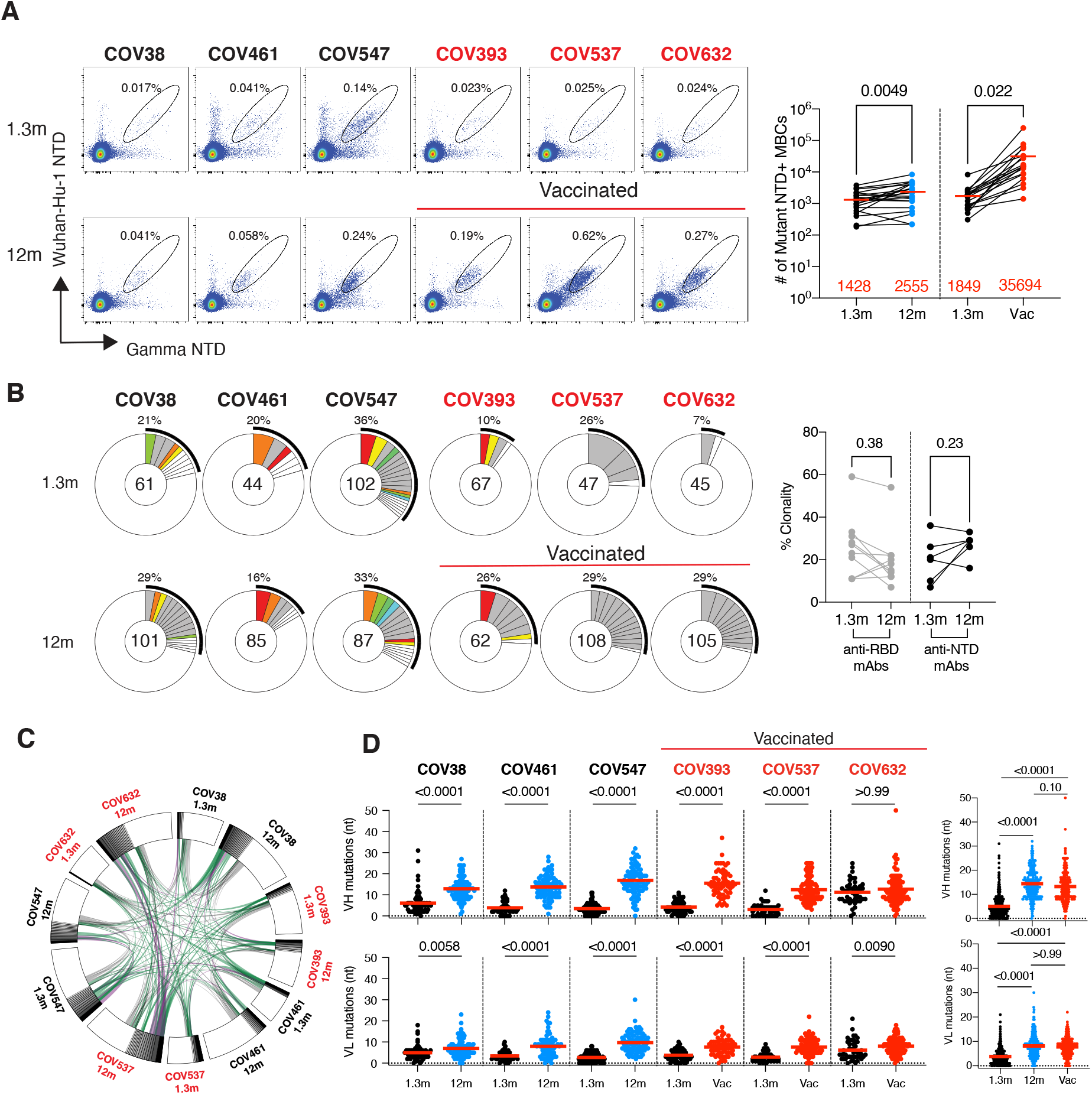
Anti-SARS-CoV-2 NTD memory B cells. **A**, Representative flow cytometry plots showing dual PE-Wuhan-Hu-1 and AlexaFluor-647-Gamma NTD binding B cells for 6 individuals (vaccinees, n=3, non-vaccinees, n=3) at 1.3 month(Robbiani et al., 2020) and 12 months(Wang et al., 2021c). Gating strategy is found in **Figure S4**. Right panel shows number of NTD/Mutant NTD positive B cells per 10 million B cells (also see in **Figure S4B and 4C**) obtained at 1.3 month and 12 months from 39 randomly selected individuals (vaccinees n=18, and non-vaccinees, n=21). Graphs showing dot plots from individuals 1.3 months after infection (black), non-vaccinated convalescents (blue) or vaccinated convalescents (red) 12m after infection. Each dot is one individual. Red horizontal bars indicate mean values. Statistical significance was determined using two-tailed Mann–Whitney U-tests. **B**, Pie charts show the distribution of antibody sequences from 6 individuals after 1.3 month (upper panel) or 12 months (lower panel). The number in the inner circle indicates the number of sequences analyzed for the individual denoted above the circle. Pie slice size is proportional to the number of clonally related sequences. The black outline indicates the frequency of clonally expanded sequences detected. Colored slices indicate persisting clones (same IGHV and IGLV genes, with highly similar CDR3s) found at both timepoints in the same patient. Grey slices indicate clones unique to the timepoint. The percentage of BCR clonality from each individual was summarized in the right panel. Statistical significance was determined using two tailed-Wilcoxon matched-pairs signed rank tests. **C**, Circus plot depicts the relationship between antibodies that share V and J gene segment sequences at both IGH and IGL. Purple, green, and grey lines connect related clones, clones and singles, and singles to each other, respectively. Vaccinees are marked in red. **D**, Number of somatic nucleotide mutations in the IGVH and IGVL in antibodies obtained after 1.3 or 12 months from 6 donors. Right panel showing dot plots from individuals 1.3 months after infection (black), non-vaccinated convalescents (blue) or vaccinated convalescents (red) 12m after infection. Red horizontal bars indicate mean values. Statistical significance was determined using Kruskal Wallis test with subsequent Dunn’s multiple comparisons.

Expanded clones of memory B cells were found at both time points. On average expanded clones accounted for 22% and 27% of all antibodies at the 1.3- and 12-month time points respectively, and 30 out of 90 such clones were conserved between time points (Figure 1B and Table S2). Thus, most of the clones were unique to one of the two time points indicating that the antibody response continued to evolve with persisting clonal expansion. B cell clonal evolution is associated with accumulation of somatic hypermutation (Elsner and Shlomchik, 2020; Victora and Nussenzweig, 2012). Consistent with the notion that the anti-NTD antibody response continues to evolve there was a significant increase in Ig heavy and/or light chain somatic mutation between the 2 time points in all 6 individuals examined (Figure 1D). CDR3 length was unchanged overtime and hydrophobicity was slightly lower for anti-NTD antibodies than control (Figure S6).

We expressed 275 antibodies from the 6 individuals (124 and 151 from 1.3- and 12-month timepoint, respectively) including: 1) 158 that were randomly selected from those that appeared only once evenly divided between 1.3- and 12-months; 2) 29 that appeared as expanded clones that were conserved at the two time points; 3) 16 and 43 newly arising expanded clones present at either the 1.3- or 12-month time points (Table S3). Each of the antibodies was tested for binding to Wuhan-Hu-1, Delta, Gamma, and Omicron NTDs by ELISA (Figure 2A). 97% of the antibodies cloned from the 1.3-month time point bound to the Gamma NTD, and 82%, 69% and 52% to Wuhan-Hu-1, Delta, and Omicron respectively. The fraction of binding antibodies to Wuhan-Hu-1 and Delta NTD improved significantly after 12 months (Figures 2A and S7A). In addition, the geometric mean ELISA half-maximal concentration (EC_50_) decreased significantly for all the variant NTDs after 12 months suggesting an increase in overall binding affinity over time (Figure 2A and Table S3).

**Figure 2.**
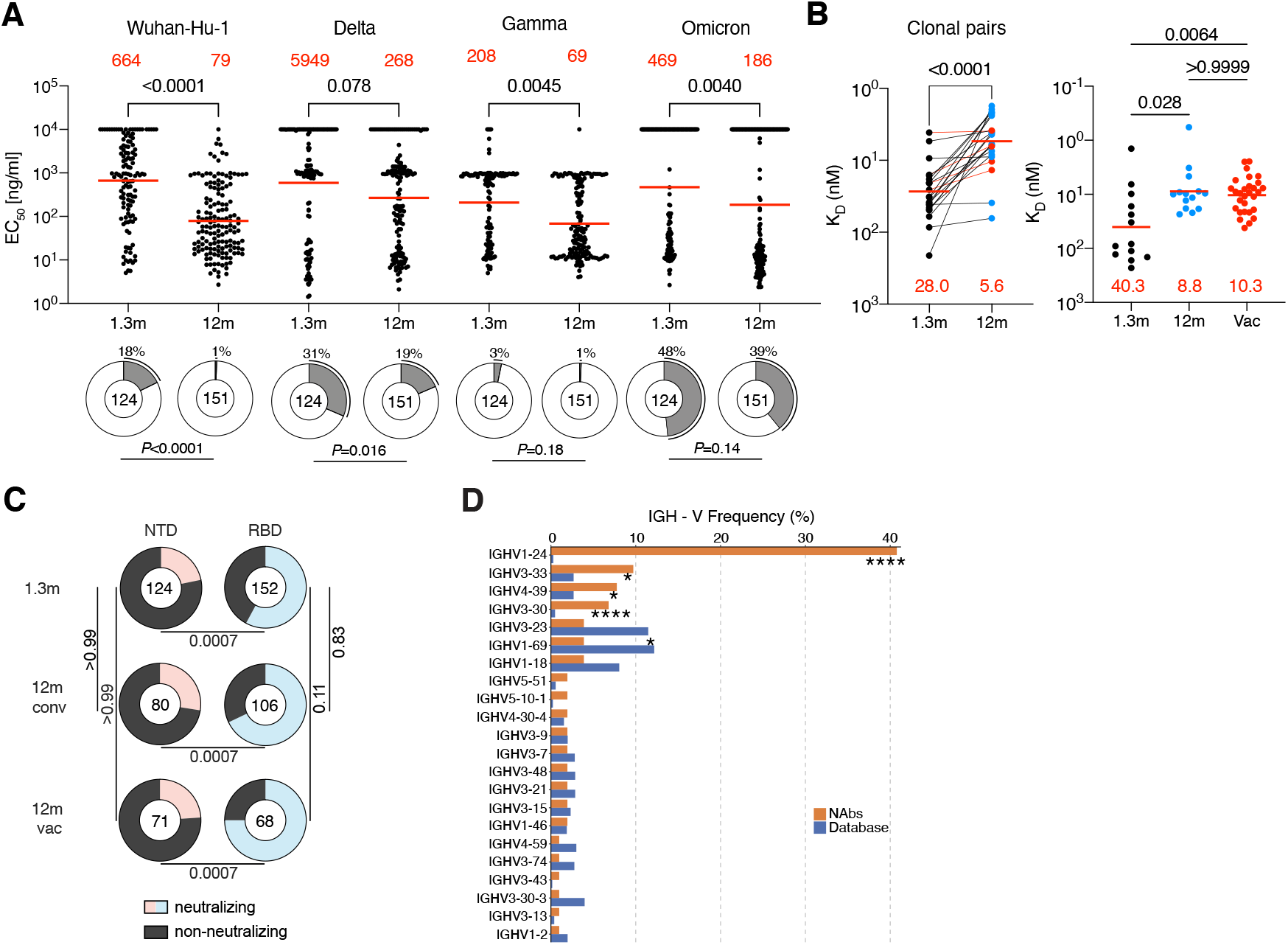
Binding and neutralizing activity of anti-SARS-CoV-2 NTD antibodies. **A**, Dot plots show EC_50_s against SARS-CoV-2 Wuhan-Hu-1-, Delta-, Gamma-, or Omicron-NTD for mAbs isolated from convalescents individuals at 1.3- and 12-months after infection. Each dot represents an individual antibody. Red horizontal bars indicate geometric mean values. Statistical significance was determined through the two-sided Kruskal Wallis test with subsequent Dunn’s multiple comparisons. Pie charts illustrate the fraction of binders (EC_50_ 0-10000 ng/ml, white slices), and non-binders (EC_50_ = 10000 ng/ml, grey slices). Inner circle shows the number of antibodies tested per group. The black outline indicates the percentage of non-binders. Statistical significance was determined with two-sided Fisher’s exact test. **B**, Graphs depict affinity measurements for mAbs obtained 1.3- and 12-months after infection. Left panel shows dissociation constants K_D_ values for 22 clonally paired antibodies. Right panel shows K_D_ values for randomly selected antibodies. Antibodies from individuals 1.3 months after infection are in black, from convalescents 12 m after infection non-vaccinated are in blue or vaccinated in red. Statistical significance was determined with two-sided Kruskal–Wallis test with subsequent Dunn’s multiple comparisons. Horizontal bars indicate geometric mean values. Statistical significance was determined using Wilcoxon matched-pairs signed rank tests. **C**, Pie charts illustrate the fraction of neutralizing (colored slices) and non-neutralizing (IC_50_ > 1000 ng/ml, grey slices) anti-NTD (left) and anti-RBD (Wang et al., 2021c) (right) monoclonal antibodies, inner circle shows the number of antibodies tested per group. Statistical significance was determined with Fisher’s exact test with subsequent Bonferroni-Dunn correction. **D**, Graph shows comparison of the frequency distributions of human IGH V genes of anti-SARS-CoV-2 NTD neutralizing antibodies from donors at 1.3 month (Robbiani et al., 2020) and 12 months(Wang et al., 2021c) after infection. Statistical significance was determined by two-sided binomial test.

To determine whether antibody affinity increased between the 2 time points we performed biolayer interferometry experiments (Figures 2A and S7B). Between 1.3- and 12-months affinity to the Wuhan-Hu-1 NTD increased among clonal pairs (28.0nM to 5.6nM) and unique antibodies (40.3nM to 8.8nM in unvaccinated or to 10.3nM in vaccinated individuals) (Figure 2B). We conclude that anti-NTD antibodies evolve to higher affinity during the 12 months following infection irrespective of subsequent vaccination.

### Anti-NTD Antibody Neutralizing Activity

Antibodies that showed binding to NTD by ELISA were tested for neutralizing activity in a SARS-CoV-2 neutralization assay using pseudotyped viruses encoding Wuhan-Hu-1, Gamma and PMS20 S proteins (Table S3). Among 275 NTD-binding antibodies, 103 neutralized at least one of the pseudoviruses with an IC_50_ of less than 1000 ng/ml (Table S3). The fraction of Wuhan-Hu-1-neutralizers among the anti-NTD antibodies obtained from convalescent individuals was 22%, 28%, and 24% at 1.3-, 12-month and 12-month vaccinated individuals, respectively. In contrast, the overall neutralizing activity among anti-RBD antibodies obtained from the same time points from this cohort was significantly higher in all cases (58%, 68%, 75% respectively) (Figure 2C, (Robbiani et al., 2020; Wang et al., 2021c)). Anti-NTD neutralizing antibodies were significantly enriched in IGVH1-24, IGVH3-33 and IGVH3-30 all of which are enriched among neutralizing antibodies that target the NTD supersite (Figure 2D and Table S4, (Amanat et al., 2021; Cerutti et al., 2021b; Chi et al., 2020; Dussupt et al., 2021; Haslwanter et al., 2021; Li et al., 2021; Liu et al., 2021; McCallum et al., 2021a; Planas et al., 2021b; Suryadevara et al., 2021b; Voss et al., 2021)). IGVH1-24 was frequently associated with IGVL1-51 which is also over-represented among neutralizers compared to the database (Table S4). The neutralizers obtained by this method were also enriched in VH4-39 which has not been associated with NTD supersite neutralizing activity (Figure 2D).

Among the antibodies tested the geometric mean IC_50_ against Wuhan-Hu-1, Gamma, PMS20 was similar at 1.3- and 12-months (Figures 3A and S8A, Table S3). Vaccination was associated with increased neutralizing activity at 12 months against Gamma but not the other 2 pseudoviruses tested (Figure 3A). There was also no change in geometric mean IC_50_ when only the conserved clones were considered (Figure S8B). Therefore, there is no general increase in anti-NTD antibody neutralizing activity over time despite the relative increase in affinity.

**Figure 3.**
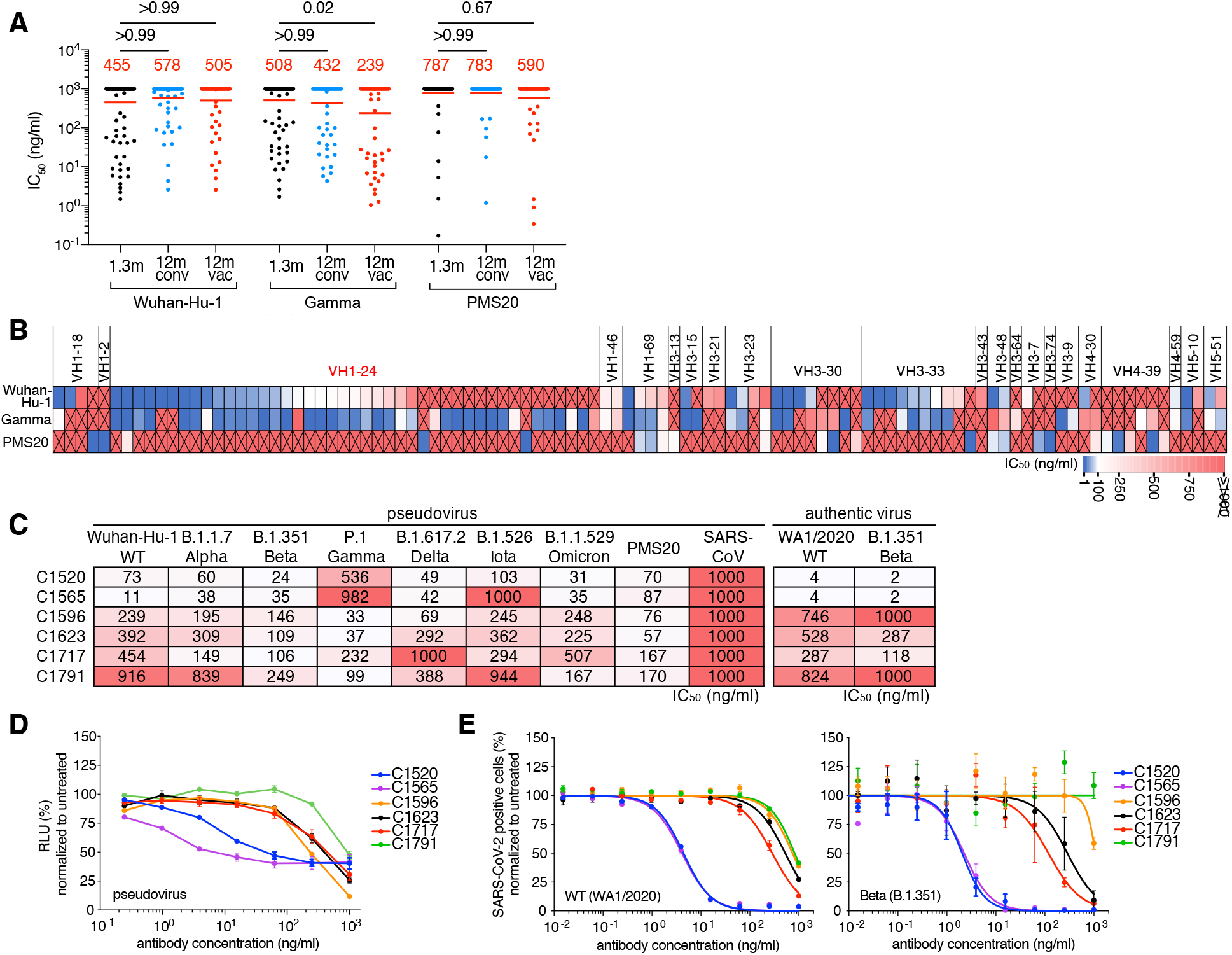
Neutralizing breath of anti-SARS-CoV-2 NTD antibodies. **A,** Graph shows anti-SARS-CoV-2 neutralizing activity of monoclonal antibodies measured by a SARS-CoV-2 pseudovirus neutralization assay(Robbiani et al., 2020; Schmidt et al., 2020). Half-maximal inhibitory concentration (IC_50_) values for antibodies isolated at 1.3 and 12 months after infection in non-vaccinated (1.3m/12m) and convalescent vaccinated (12m vac) participants. IC_50_s against Wuhan-Hu-1, gamma and PMS20(Schmidt et al., 2021c) SARS-CoV-2. Spike plasmids are based on Wuhan-Hu-1 strain containing the R683G substitution. Each dot represents one antibody from individuals 1.3 months after infection (black), non-vaccinated convalescents (blue) or vaccinated convalescents (red) 12m after infection. Statistical significance was determined by two-sided Kruskal Wallis test with subsequent Dunn’s multiple comparisons. Horizontal bars and red numbers indicate geometric mean values. **B**, Heatmap showing neutralizing activity and VH groups of 103 neutralizing antibodies against Wuhan-Hu-1, Gamma and PMS20 SARS-CoV-2 pseudovirus. IC_50_ values are indicated by color (from blue to red, most potent to least potent) with non-neutralizing antibodies (IC50 values of >1000 ng/ml) marked by crossed red tiles. **C**, IC_50_ values for n=6 broadly neutralizing NTD antibodies against the indicated variant SARS-CoV-2 pseudoviruses, PMS20(Schmidt et al., 2021c) and SARS-CoV pseudovirus(Muecksch et al., 2021) (left) as well as WT (WA1/2020)(Robbiani et al., 2020) and Beta (B.1.351) authentic virus. **D**, Neutralization curves for the antibodies in **C,** against Wuhan-Hu-1 pseudovirus. **E**, Neutralization curves for the antibodies in **D,** against WT (WA1/2020) and Beta (B.1.351) authentic virus.

Although the neutralizing activity of the NTD antibodies was generally lower than RBD antibodies cloned from the same time points from this cohort (Figures 3A and S8A, (Robbiani et al., 2020; Wang et al., 2021c)), some anti-NTD antibodies were very potent with IC_50_ values as low as 0.17 nanograms per milliliter (Figure 3A). Of the 103 neutralizing anti-NTD antibodies with demonstrable neutralizing activity 14 were specific for Wuhan-Hu-1, 20 were limited to Gamma, and 13 were PMS20-specific (Figure 3B). The remaining 56 antibodies neutralized 2 or more viruses (Figure 3B). Antibodies targeting the NTD supersite are enriched in VH1-24, VH3-30 and VH3-33 and these 3 VH genes account for 59 of the 103 antibodies tested (Figures 3B and S8C, Table S4). Notably, despite its mutations in the NTD supersite, 24 antibodies neutralized PMS20 and 6 neutralized all 3 viruses suggesting that some of these antibodies might bind to epitopes outside of the supersite.

To document the neutralizing breadth of the 6 broadest antibodies, we tested them against viruses pseudotyped with SARS-CoV-2 Alpha, Beta, Delta, Iota and Omicron and SARS-CoV S proteins (Figure 3C). Although none of the antibodies neutralized SARS-CoV, 4 of the 6 antibodies neutralized all strains tested albeit at relatively high neutralizing concentrations. However, pseudovirus neutralization was incomplete even at very high antibody concentrations for 2 of the more potent antibodies tested (Figure 3D).

To determine whether intact virus neutralization resembles pseudovirus neutralization we performed microneutralization experiments using authentic SARS-CoV-2-WA1/2020 (Robbiani et al., 2020) and -Beta. In contrast to the pseudovirus, the two antibodies that showed the most incomplete neutralization profiles against pseudovirus, C1520 and C1565, reached complete neutralization and were exquisitely potent with IC_50_s in the low nanogram per milliliter range against both strains (Figures 3C and 3E). We conclude that some naturally arising memory anti-NTD antibodies produced in response to Wuhan-Hu-1 infection and immunization are insensitive to the mutations found in Omicron and other variants of concern.

To determine whether our collection of anti-NTD neutralizing antibodies target overlapping epitopes we performed biolayer interferometry competition experiments (Figures 4A and 4B). Among the 43 antibodies with the highest neutralizing activity tested there were 6 discernible complementation groups. Groups I and II were overlapping and highly enriched in VH3-30, VH3-33 and VH1-24 respectively accounting for nearly 90% of the antibodies in these 2 groups (Figure 4B). These antibodies neutralized either or both Wuhan-Hu-1 and Gamma, but none neutralized PMS20 or omicron that are mutated in the NTD supersite. Thus, these 2 groups of antibodies appear to target the previously defined supersite on NTD (Amanat et al., 2021; Cerutti et al., 2021b; Chi et al., 2020; Dussupt et al., 2021; Haslwanter et al., 2021; Li et al., 2021; Liu et al., 2021; McCallum et al., 2021a; Planas et al., 2021b; Suryadevara et al., 2021b; Voss et al., 2021). Group III and IV are also overlapping but in contrast, groups III and IV only 3 of the 19 neutralize Wuhan-Hu-1 and those that do are broad. Notably, the remaining antibodies fail to measurably neutralize Wuhan-Hu-1 but neutralize PMS20 and/or Omicron (Figure 4B and Table S4). Group V contains 4 members, 3 of which are broad. The final group VI contains 2 members, C1621 only neutralizes Wuhan-Hu-1, and C1554 neutralized broadly but not potently. The broadest neutralizing anti-NTD antibodies appear to recognize sites outside of the supersite and are not dominated by VH1-24 and VH3-30/3-33. Altogether 16 out of the 43 anti-NTD neutralizing antibodies tested neutralized PMS20 and/or Omicron but not Wuhan-Hu-1. Thus, the B cell memory compartment produced in response to infection with Wuhan-Hu-1 contains antibodies that bind to this strain with high affinity, but do not neutralize it and instead neutralize PMS20 and/or Omicron.

**Figure 4.**
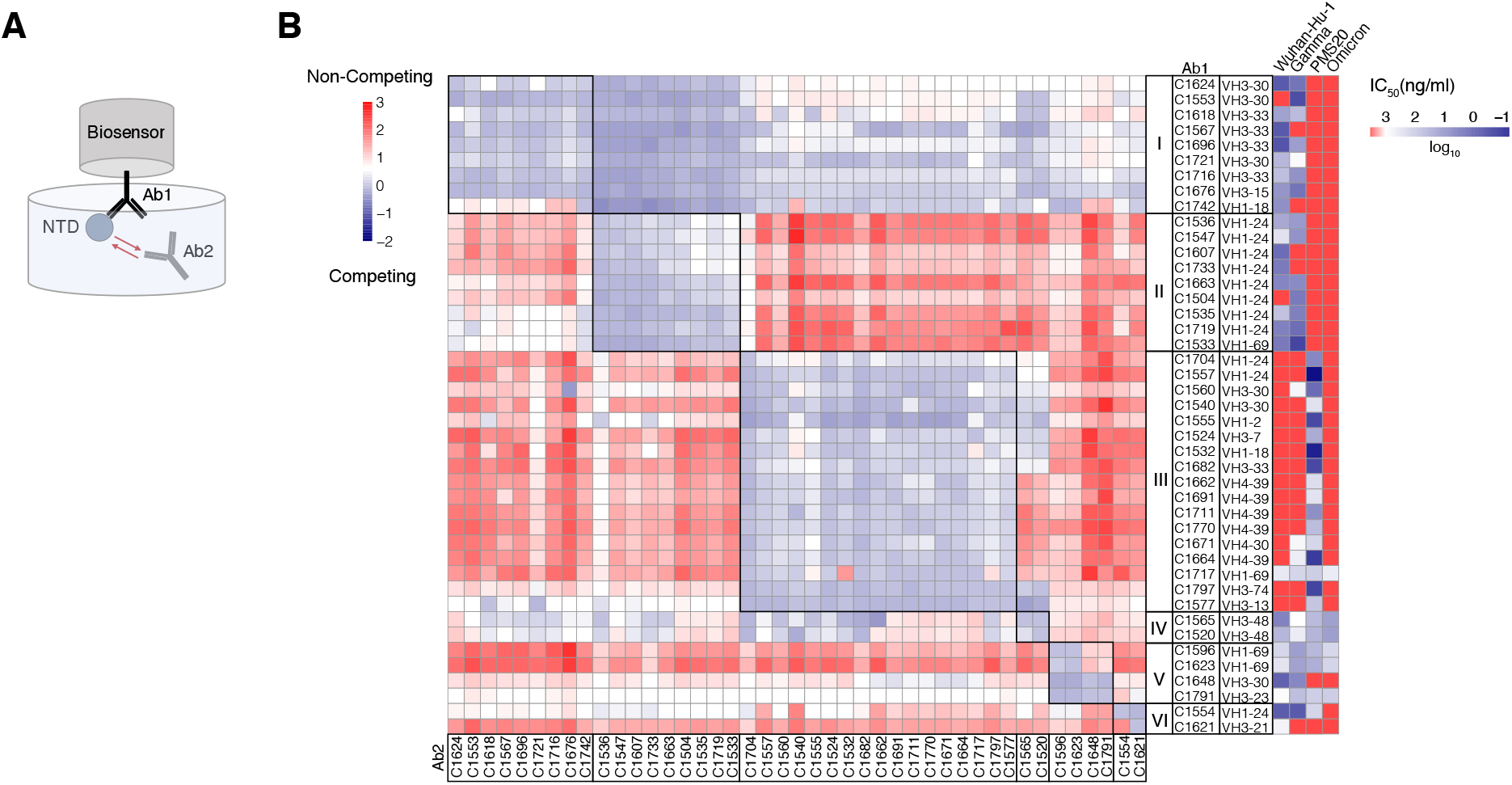
Neutralizing epitope mapping of anti-SARS-CoV-2 NTD antibodies. **A**, Diagram of the biolayer interferometry experiment. **B**, Biolayer interferometry results presented as a heat-map of relative inhibition of Ab2 binding to the preformed Ab1-NTD complexes (from red to blue, non-competing to competing). Values are normalized through the subtraction of the autologous antibody control. VH gene usage and IC_50_ values of Wuhan-Hu-1, Gamma, PMS20 and Omicron (heat map, from blue to red, most potent to least potent) for each antibody are shown. The average of two experiments is shown.

### Neutralizing epitopes on NTD outside the supersite

To delineate the structural basis for broad-recognition of NTD-directed antibodies, we determined structures of WT SARS-CoV-2 S 6P(Hsieh et al., 2020) bound to Fab fragments of C1717 (group III), C1520 (group IV), and C1791 (group V) using single-particle cryo-electron microscopy (cryo-EM) (Figure S9 and Table S5). Global refinements yielded maps at 2.8Å (C1520-S), 3.5Å (C1717-S), and 4.5Å (C1791-S) resolutions, revealing Fab fragments bound to NTD epitopes on all three protomers within a trimer irrespective of ‘up’/‘down’ RBD conformations for all three Fab-S complexes. C1520 and C1791 Fabs recognize epitopes on opposite faces of the NTD, with binding poses orthogonal to the site i antigenic supersite and distinct from C1717 pose, consistent with BLI mapping data (Figures 4b and S9J). A 4.5Å resolution structure of antibody C1791 (*VH3-23*01/VK1-17*01*) bound to a S trimer revealed a glycopeptidic NTD epitope wedged between the N61_NTD_- and N234_NTD_-glycans, engaging several N-terminal regions including the N1-, N2- and b8-b9 hairpin loops (Figure S9K). The binding pose of C1791 is similar to the cross-reactive antibody S2L20, which was shown to maintain binding against single NTD mutations and several VOCs but was non-neutralizing(McCallum et al., 2021a; McCallum et al., 2021b).

Antibody C1520 (*VH3-48*03/VL4-68*02*) recognizes the NTD b-sandwich fold, with a distinct pose relative to similar cross-reactive NTD mAbs (Figures 5A, 5B and S10A)(Cerutti et al., 2021a; McCallum et al., 2021b). The C1520 epitope comprises residues along the supersite b-hairpin (residues 152-158), the b8-strand (residue 97-102), N4-loop (residues 178-188), and N-linked glycans at positions N122 and N149 (Figure 5C). Targeting of the NTD epitope was driven primarily by heavy chain contacts (the buried surface area (BSA) of NTD epitope on the C1520 HC represented ∼915Å^2^ of ∼1150Å^2^ total BSA), mediated by the 20-residue long CDRH3 that contributed 55% of the antibody paratope (Figures 5C and 5D). Previous studies have shown that CDRH3 loops of antibodies targeting this face of the NTD disrupt a conserved binding pocket that accommodates hydrophobic ligands, including polysorbate 80 and biliverdin, which have the potential to prevent binding of supersite antibodies(Cerutti et al., 2021a; Rosa et al., 2021). The 20-residue long CDRH3 of C1520 displaces the supersite b-hairpin and N4 loops, which acts as a gate for the hydrophobic pocket (Figure 5E), in a manner similar to antibody P008-056 ( Figure S10B). This contrasts Ab5-7 that directly buries the tip of its CDRH3 into the hydrophobic pocket, and S2X303 that maintains a closed gate and partially-overlaps with the supersite b-hairpin ( Figure S10B). Thus, C1520’s increased cross-reactivity and neutralization breadth relative to Ab5-7 and S2X303 is likely mediated by displacement of the supersite b-hairpin and N4-loops, which harbor escape mutations found in several SARS-CoV-2 VOCs and can undergo structural remodeling to escape antibody pressure (Figure 5E)(McCallum et al., 2021b).

**Figure 5.**
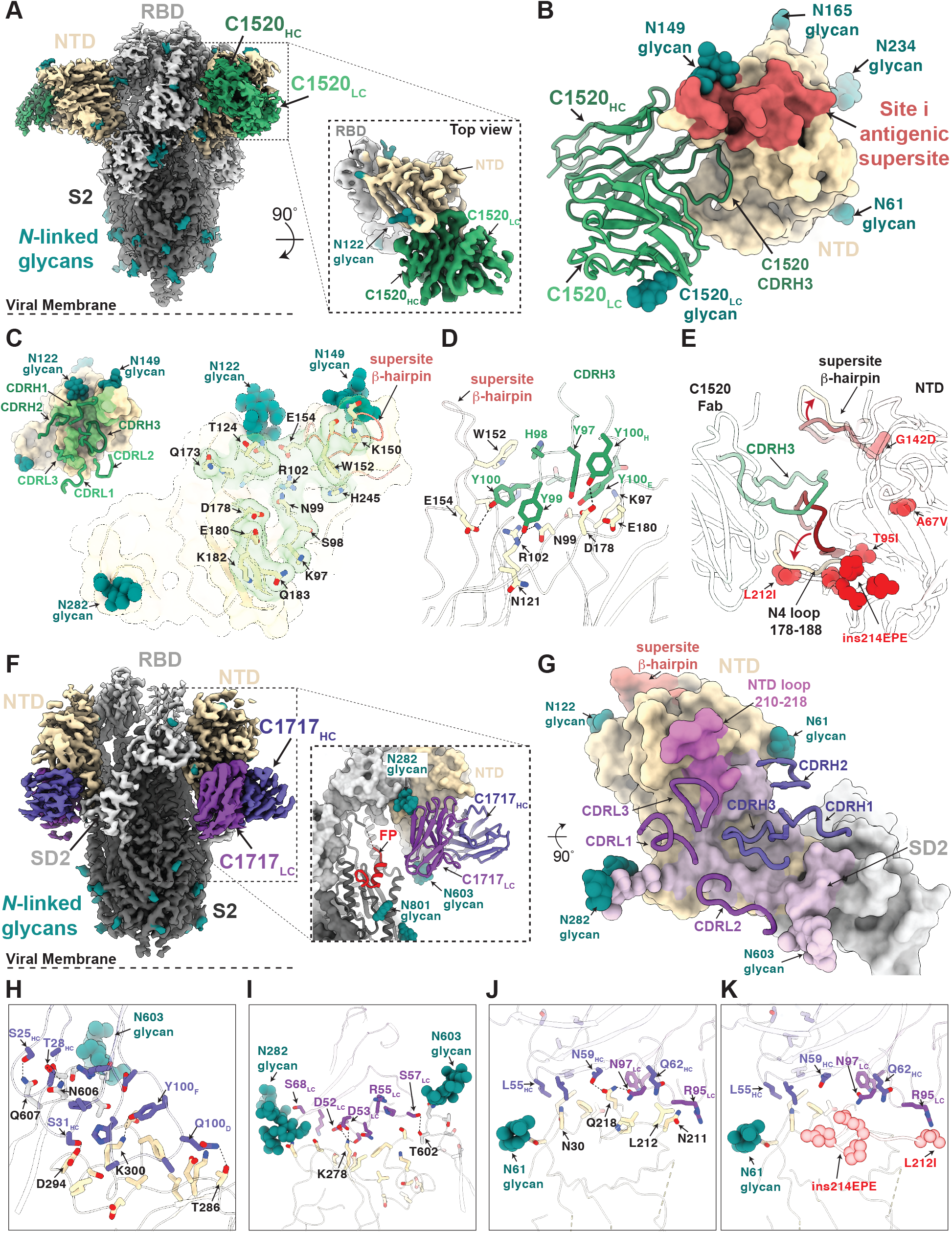
Cryo-EM structures of C1520 and C1717 bound to the SARS-CoV-2 S 6P ectodomain. **A,** 2.8Å cryo-EM density for C1520-S 6P trimer complex. Inset: 3.1Å locally-refined cryo-EM density for C1520 variable domains (shades of green) bound to NTD (wheat). **B,** Close-up view of C1520 variable domains (green ribbon) binding to NTD (surface rendering, wheat). Site i antigenic supersite is highlighted on NTD for reference (salmon). **C,** Surface rendering of C1520 epitope (green) with CDR loops shown (inset). NTD epitope residues (defined as residues containing atom(s) within 4Å of a Fab atom) are shown as sticks. **D,** CDRH3-mediated contacts on the NTD. Potential hydrogen bonds are shown as black dashed lines. **E,** Overlay of WT (WA1/2020) NTD (this study) and unliganded Omicron NTD (PDB 7TB4) with mutations and loop conformational changes highlighted in red. **F,** 3.5Å cryo-EM density for C1717-S 6P trimer complex. Inset: C1717 variable domains (shades of purple) binding to a S1 protomer (surface rendering). The fusion peptide (FP, red) in the S2 domain is shown. **G,** Surface rendering of C1717 epitope (thistle) with C1717 CDR loops shown as ribbon. **H-K,** Residue-level contacts between C1717 (purple), NTD (wheat) and the SD2 domain (gray). Omicron mutations in the N5 loop are shown as red spheres in panel **K**.

We next investigated the group III antibody, C1717, that showed neutralization breadth against SARS-CoV-2 VOCs and distinct binding properties from group IV and V antibodies (Figures 3C, 4B and S9J). Antibody C1717 (*VH1-69*04/VL1-44*01*) recognizes the viral membrane proximal side of the NTD (Figure 5F), similar to S2M24(McCallum et al., 2021a), DH1052(Li et al., 2021), and polyclonal Fabs from donor COV57(Barnes et al., 2020b). However, C1717’s pose is unique and represents a new antigenic site that recognizes a glycopeptide epitope flanked by the N282_NTD_- and N603_SD2_-glycans and is positioned in close proximity (<12 Å) to the S2-fusion peptide region (Figures 5F and 5G). All six CDR loops contribute to an epitope that spans both the NTD and SD2 regions (residues 600-606) with a spike epitope BSA of ∼1325 Å^2^ (Figure 5G). CDRH1 and CDRH3 loops mediate extensive hydrogen bond and van der Waals contacts with the C-terminus of the NTD (residues 286-303) and SD2 loop (residues 600-606), whereas CDRH2 engages N-terminal regions (residues 27-32, 57-60) and NTD loop residues 210-218 (Figures 5H-5K). C1717 light chain CDRL2 and CDRL3 loops contact the C-terminus of the NTD (residues 286-303) and the NTD loop 210-218, respectively (Figures 5I and 5J). In addition, light chain FWR3 buries against the N282_NTD_-glycan, a complex-type N-glycan(Wang et al., 2021a) that wedges against the SD1 domain of the adjacent protomer (Figures 5F-G, I). Modeling of Omicron mutations found in NTD loop 210-218 shows accommodation of the 214-insertion and potential establishment of backbone contacts with the NTD loop and R95_LC_ in CDRL3 (Figure 5K). Moreover, stabilization of the N282_NTD_-glycan against the adjacent protomer and the proximity of the C1717 LC to the S2 fusion machinery potentially contributes to C1717’s neutralization mechanism by preventing access to the S2’ cleavage site or destabilization of S1. Such a mechanism could explain the lack of Delta VOC neutralization, as this variant has been shown to have enhanced cell-cell fusion activity relative to all other VOCs(McCallum et al., 2021b; Zhang et al., 2021).

## Discussion

In animal models, passive antibody infusion early in the course of infection accelerates viral clearance and protects against disease (Hansen et al., 2020; Hassan et al., 2020; Rogers et al., 2020; Schmitz et al., 2021; Tortorici et al., 2020; Zost et al., 2020a). In humans, rapid development of neutralizing antibodies to SARS-CoV-2 is associated with better clinical outcomes (Khoury et al., 2021). Finally, early passive transfer of neutralizing antibodies to high-risk individuals alters the course of the infection and can prevent hospitalization and death (Dougan et al., 2021; Gupta et al., 2021; Weinreich et al., 2021). Thus, antibodies play an essential role in both protection against SARS-CoV-2 infection and serious disease.

The RBD and the NTD are the two dominant targets of neutralizing antibodies on the SARS-CoV-2 S protein. When the memory B cell compartment is probed using intact S the majority of the neutralizing antibodies obtained target the RBD and only a small number that are usually less potent or broad target the NTD (Kreer et al., 2020; Liu et al., 2020; Zost et al., 2020b). Among the neutralizers that recognize the NTD, the majority target the site i supersite and these antibodies are enriched in IGVH3-33 and IGVH1-24 (Amanat et al., 2021; Cerutti et al., 2021b; Chi et al., 2020; Dussupt et al., 2021; Haslwanter et al., 2021; Li et al., 2021; Liu et al., 2021; McCallum et al., 2021a; Planas et al., 2021b; Suryadevara et al., 2021b; Voss et al., 2021). The few existing exceptions target a region orthogonal to the supersite featuring a pocket that accommodates hydrophobic ligands, such as polysorbate 80 and biliverdin (Cerutti et al., 2021a; Rosa et al., 2021). Focusing on the NTD and using complementary probes revealed 6 partially overlapping complementary groups of NTD neutralizing antibodies.

Groups I and II are partially overlapping and appear to target the supersite on the target cell facing surface of the NTD. Group I is enriched in IGVH3-33 and IGVH3-30, while group II is enriched in IGVH1-24. Together, these two groups make up 42% of all the neutralizers characterized, none of which are broad or able to neutralize Omicron or PMS20 that carry extensive site i supersite mutations. Groups I and II differ in their ability to inhibit each other’s binding to NTD likely because they assume different binding poses. Group III is enriched in IGVH4-39 and is unusual in that all the antibodies in this group neutralize the synthetic PMS20 variant, but only 24% neutralize Gamma, and only 1 of 18 tested is broad. The antibodies in this group target a novel site of neutralization on the membrane proximal side of the NTD, bridging the SD2 domain on the same protomer, and potentially infringe on the S2 domain to impede spike function in viral-host membrane fusion(Suryadevara et al., 2021a). Groups IV, V and VI comprise only 19% of all the NTD neutralizers tested but are highly enriched in the broad neutralizers, which have the potential to provide universal protection against SARS-CoV-2 variants infection. Group IV and V antibodies target opposite sides of the NTD orthogonal to the site i supersite, recognizing epitopes that are non- or only partially-overlapping with VOC mutations and are not significantly impacted by conformational changes in the supersite b-hairpin(McCallum et al., 2021b). While structurally unique in their binding pose, antibodies Group IV and V antibodies likely fail to inhibit ACE2 interactions suggesting a potent neutralization mechanism that stabilizes the prefusion spike glycoprotein. Such mechanisms have been suggested for similar NTD-directed antibodies(Cerutti et al., 2021a; McCallum et al., 2021a; Suryadevara et al., 2021b), and remains to be explored.

Although a great number of SARS-CoV-2 neutralizing antibodies have been identified, Omicron escapes the majority of them, including most antibodies in clinical use (Cameroni et al., 2021; Cao et al., 2021; Planas et al., 2021a). Identifying new epitopes that are targeted by the broad neutralizers and are conserved among SARS-CoV-2 variants would be ideal targets for future development of broadly active sarbecovirus vaccines and antibody drugs to overcome antigenic drift.

Immune responses selectively expand two independent groups of B lymphocytes, plasma cells and memory B cells. Plasma cells are end stage cells that are selected based on affinity. They are home to the bone marrow where they reside for variable periods of time and secrete specific antibodies found in plasma. An initial contraction in the number of these cells in the first 6 months after SARS-CoV-2 infection or vaccination likely accounts for the decrease in plasma neutralizing activity and subsequent loss of protection from infection(Cho et al., 2021; Gaebler et al., 2021).

Memory cells are found in circulation and in lymphoid organs where they are relatively quiescent. B cells selected into this compartment are far more diverse than those that become plasma cells and include cells with somatically mutated receptors that have variable affinities for the immunogen (Viant et al., 2020; Viant et al., 2021). This includes cells producing antibodies with relatively high affinities that are highly specific for the antigen as well as cells producing antibodies with affinity for the initial immunogen and the ability to bind to closely related antigens. Production of a pool of long lived but quiescent memory cells expressing a diverse collection of closely related antibodies, some of which can recognize pathogen variants, favors rapid secondary responses upon SARS-CoV-2 variant exposure in pre-exposed and vaccinated individuals.

SARS-CoV-2 infection or vaccination with Wuhan-Hu-1 produces plasma antibody responses that are relatively specific for Wuhan-Hu-1 and are sub-optimally protective against infection with variant strains such as Omicron. In contrast, memory B cell populations produced in response to infection or vaccination are sufficiently diversified to contain high affinity antibodies that can potently neutralize numerous different variants including Omicron. This diverse collection of memory cells and their cognate memory T cells are likely to make a significant contribution to the generally milder course of infection in individuals that have been infected and/or vaccinated with SARS-CoV-2.

## Acknowledgements

We thank all study participants who devoted time to our research, The Rockefeller University Hospital nursing staff and Clinical Research Support Office. Thanks all members of the M.C.N. laboratory for helpful discussions, Maša Jankovic for laboratory support and Kristie Gordon for technical assistance with cell-sorting experiments. We thank the laboratory of Dr. Pamela Bjorkman at the California Institute of Technology for SARS-CoV-2 spike expression constructs. Cryo-EM data for this work was collected at the Stanford-SLAC cryo-EM center with support from Dr. Elizabeth Montabana and the Evelyn Gruss Lipper cryo-EM resource center (Rockefeller University) with support from Mark Ebrahim, Johanna Sotiris and Honkit Ng. This work was supported by NIH grant P01-AI138398-S1 (M.C.N.) and 2U19AI111825 (M.C.N.). R37-AI64003 to P.D.B.; R01AI78788 to T.H.; C.O.B. is supported by the Howard Hughes Medical Institute Hanna Gray Fellowship and is a Chan Zuckerberg Biohub investigator. C.G. was supported by the Robert S. Wennett Post-Doctoral Fellowship and the Shapiro-Silverberg Fund for the Advancement of Translational Research. C.G. and Z.W. were supported in part by the National Center for Advancing Translational Sciences (National Institutes of Health Clinical and Translational Science Award program, grant UL1 TR001866). P.D.B. and M.C.N. are Howard Hughes Medical Institute Investigators.

## Author Contributions

Z.W., F.M., P.D.B., T.H., C.O.B and M.C.N. conceived, designed and analyzed the experiments. M. Caskey and C.G. designed clinical protocols. Z.W., F.M., A.C., H.- H.H., S.Z., F.S., J. D.S, E.B., T.B.T., R.R., K.Y., Y.L., T.C., and C.O.B., carried out experiments. M. Cipolla., B.J. and A.G., produced antibodies. M.T., K.G.M., J. D., A.K., C.G. and M. Caskey recruited participants, executed clinical protocols and processed samples. T.Y.O. and V.R. performed bioinformatic analysis. Z.W., F.M., P.D.B., T.H., C.O.B and M.C.N. wrote the manuscript with input from all co-authors.

## METHODS

### Study participants

Samples were obtained from 62 individuals under a study protocol approved by the Rockefeller University in New York from April 1, 2020 to March 26, 2021 as described in (Robbiani et al., 2020; Wang et al., 2021c). All participants provided written informed consent before participation in the study, and the study was conducted in accordance with Good Clinical Practice. For detailed participant characteristics see Table S1.

### Blood samples processing and storage

Peripheral Blood Mononuclear Cells (PBMCs) obtained from samples collected at Rockefeller University were purified as previously reported by gradient centrifugation and stored in liquid nitrogen in the presence of FCS and DMSO (Gaebler et al., 2021; Robbiani et al., 2020). Heparinized plasma and serum samples were aliquoted and stored at −20°C or less. Prior to experiments, aliquots of plasma samples were heat-inactivated (56°C for 1 hour) and then stored at 4°C.

### ELISAs

ELISAs(Amanat et al., 2020; Grifoni et al., 2020) to evaluate antibodies binding to SARS-CoV-2 Wuhan-Hu-1-, Delta-, Gamma and Omicron-NTD were performed by coating of high-binding 96-half-well plates (Corning 3690) with 50 μl per well of a 1μg/ml protein solution in PBS overnight at 4 °C. Plates were washed 6 times with washing buffer (1× PBS with 0.05% Tween-20 (Sigma-Aldrich)) and incubated with 170 μl per well blocking buffer (1× PBS with 2% BSA and 0.05% Tween-20 (Sigma)) for 1 h at room temperature. Immediately after blocking, monoclonal antibodies or plasma samples were added in PBS and incubated for 1 h at room temperature. Plasma samples were assayed at a 1:66 starting dilution and 7 additional threefold serial dilutions. Monoclonal antibodies were tested at 10 μg/ml starting concentration and 10 additional fourfold serial dilutions. Plates were washed 6 times with washing buffer and then incubated with anti-human IgG, IgM or IgA secondary antibody conjugated to horseradish peroxidase (HRP) (Jackson Immuno Research 109-036-088 109-035-129 and Sigma A0295) in blocking buffer at a 1:5,000 dilution (IgM and IgG) or 1:3,000 dilution (IgA). Plates were developed by addition of the HRP substrate, TMB (ThermoFisher) for 10 min (plasma samples) or 4 minutes (monoclonal antibodies), then the developing reaction was stopped by adding 50 μl 1 M H_2_SO_4_ and absorbance was measured at 450 nm with an ELISA microplate reader (FluoStar Omega, BMG Labtech) with Omega and Omega MARS software for analysis. For plasma samples, a positive control (plasma from participant COV57, diluted 66.6-fold and seven additional threefold serial dilutions in PBS) was added to every assay plate for validation. The average of its signal was used for normalization of all of the other values on the same plate with Excel software before calculating the area under the curve using Prism V9.1(GraphPad). For monoclonal antibodies, the EC_50_ was determined using four-parameter nonlinear regression (GraphPad Prism V9.1).

### Expression of NTD proteins

Mammalian expression vectors encoding the SARS-CoV-2 Wuhan-Hu-1-NTD (GenBank MN985325.1; S protein residues 14-307), or Delta, Gamma and Omicron NTD mutants with an N-terminal human IL-2 or Mu phosphatase signal peptide and a C-terminal polyhistidine tag followed by an AviTag were used to express soluble NTD proteins by transiently-transfecting Expi293F cells (GIBCO). After four days, NTD proteins were purified from the supernatants by nickel affinity and size-exclusion chromatography. Peak fractions were identified by SDS-PAGE, and fractions corresponding to monomeric NTDs were pooled and stored at 4°C.

### Cell Lines

293T cells (Homo sapiens; sex: female, embryonic kidney) obtained from the ATCC (CRL-3216) and HT1080Ace2 cl14 cells (parental HT1080: homo sapiens; sex: male, fibrosarcoma) (Schmidt et al., 2020) were cultured in Dulbecco’s Modified Eagle Medium (DMEM) supplemented with 10% fetal bovine serum (FBS) at 37 °C and 5% CO_2_. VeroE6 cells (*Chlorocebus sabaeus*; sex: female, kidney epithelial) obtained from the ATCC (CRL-1586^™^) and from Ralph Baric (University of North Carolina at Chapel Hill), and Caco-2 cells (*Homo sapiens*; sex: male, colon epithelial) obtained from the ATCC (HTB-37^™^) were cultured in DMEM supplemented with 1% nonessential amino acids (NEAA) and 10% FBS at 37 °C and 5% CO_2_. Expi293F cells (GIBCO) for protein expression were maintained at 37°C and 8% CO2 in Expi293 Expression medium (GIBCO), transfected using Expi293 Expression System Kit (GIBCO) and maintained under shaking at 130 rpm. All cell lines have been tested negative for contamination with mycoplasma.

### SARS-CoV-2 and sarbecovirus spike protein pseudotyped reporter virus

Plasmids pSARS-CoV-2-SΔ19(R683G) and pSARS-CoV-SΔ19 expressing C-terminally truncated SARS-CoV-2 and SARS-CoV spike proteins and the polymutant PMS20 spike were as described before (Schmidt et al., 2021c). A panel of plasmids expressing spike proteins from SARS-CoV-2 variants were based on pSARS-CoV-2-SΔ19(R683G) and contain the following substitutions/deletions: Alpha (B.1.1.7): ΔH69/V70, ΔY144, N501Y, A470D, D614G, P681H, T761I, S982A, D118H; Beta (B.1.351): D80A, D215G, L242H, R246I, K417N, E484K, N501Y, D614G, A701V; Gamma (P.1): L18F, T20N, P26S, D138Y, R190S, K417T, E484K, N501Y, D614G, H655Y, T1027I, V1167F; Delta (B.1.617.2): T19R, Δ156-158, L452R, T478K, D614G, P681R, D950N; Iota (B.1.526): L5F, T95I, D253G, E484K, D614G, A701V (Cho et al., 2021); Omicron (B.1.1.529) ^8^: A67V, Δ69-70, T95I, G142D, Δ143-145, Δ211, L212I, ins214EPE, G339D, S371L, S373P, S375F, K417N, N440K, G446S, S477N, T478K, E484A, Q493K, G496S, Q498R, N501Y, Y505H, T547K, D614G, H655Y, H679K, P681H, N764K, D796Y, N856K, Q954H, N969H, N969K, L981F. All SARS-CoV-2 spike proteins including variants and the polymutant spike protein PMS20 included the R683G substitution, which disrupts the furin cleavage site and generates higher titer virus stocks without significant effects on pseudotyped virus neutralization sensitivity (Schmidt et al., 2021c). SARS-CoV-2 pseudotyped particles were generated as previously described (Robbiani et al., 2020; Schmidt et al., 2020). Briefly, 293T cells were transfected with pNL4-3ΔEnv-nanoluc(Robbiani et al., 2020; Schmidt et al., 2020) and either spike plasmid. Particles were harvested 48 hpt, filtered and stored at −80C.

### Pseudotyped virus neutralization assay

Monoclonal antibodies were initially screened at a at a concentration of 1000 ng/ml to identify those that show >40% neutralization at this concentration. Antibodies were incubated with SARS-CoV-2 Wuhan-Hu-1(Robbiani et al., 2020) or SARS-CoV pseudotyped virus for 1 h at 37 °C. The mixture was subsequently incubated with HT1080Ace2 cl14 cells (Schmidt et al., 2020) for 48 h after which cells were washed with PBS and lysed with Luciferase Cell Culture Lysis 5× reagent (Promega). Nanoluc Luciferase activity in lysates was measured using the Nano-Glo Luciferase Assay System (Promega) with the Glomax Navigator (Promega). The obtained relative luminescence units were normalized to those derived from cells infected with pseudotyped virus in the absence of monoclonal antibodies. Antibodies that showed >40% neutralization at a concentration of 1000 ng/ml were subjected to further titration experiments to determine their IC_50_s. Antibodies were 4-fold serially diluted and tested against pseudoviruses as detailed above. IC_50_s were determined using four-parameter nonlinear regression (least squares regression method without weighting; constraints: top=1, bottom=0) (GraphPad Prism).

### Virus and Virus Titration

SARS-CoV-2 strains USA-WA1/2020 and the South African beta variant B.1.351 were obtained from BEI Resources (catalog no. NR-52281 and NR-54008, respectively). The original WT virus was amplified in Caco-2 cells, which were infected at a multiplicity of infection (MOI) of 0.05 plaque forming units (PFU)/cell and incubated for 6 days at 37 °C. The B.1.351 variant was amplified in VeroE6 cells obtained from the ATCC that were engineered to stably express TMPRSS2 (VeroE6_TMPRSS2_). VeroE6_TMPRSS2_ cells were infected at a MOI = 0.1 PFU/cell and incubated for 4 days at 33 °C. Virus-containing supernatants were subsequently harvested, clarified by centrifugation (3,000 *g* × 10 min), filtered using a disposable vacuum filter system with a 0.22 μm membrane and stored at −80 °C. Virus stock titers were measured by standard plaque assay (PA) on VeroE6 cells obtained from Ralph Baric (referred to as VeroE6_UNC_). Briefly, 500 µL of serial 10-fold virus dilutions in Opti-MEM were used to infect 4×10^5^ cells seeded the day prior into wells of a 6-well plate. After 1.5 h adsorption, the virus inoculum was removed, and cells were overlayed with DMEM containing 10% FBS with 1.2% microcrystalline cellulose (Avicel). Cells were incubated for 4 days at 33 °C, followed by fixation with 7% formaldehyde and crystal violet staining for plaque enumeration. All SARS-CoV-2 experiments were performed in a biosafety level 3 laboratory.

To confirm virus identity and evaluate for unwanted mutations that were acquired during the amplification process, RNA from virus stocks was purified using TRIzol Reagent (ThermoFisher Scientific, catalog no. 15596026). Brief, 200 μL of each virus stock was added to 800 μL TRIzol Reagent, followed by 200 μL chloroform, which was then centrifuged at 12,000 *g* x 5 min. The upper aqueous phase was moved to a new tube, mixed with an equal volume of isopropanol, and then added to a RNeasy Mini Kit column (QIAGEN, catalog no. 74014) to be further purified following the manufacturer’s instructions. Viral stocks were subsequently confirmed via next generation sequencing using libraries for Illumina MiSeq.

### Microscopy-Based Neutralization Assay

The day prior to infection VeroE6_UNC_ cells were seeded at 1×10^4^ cells/well into 96-well plates. Antibodies were serially diluted (4-fold) in BA-1, consisting of medium 199 (Lonza, Inc.) supplemented with 1% bovine serum albumin (BSA) and 1x penicillin/streptomycin. Next, the diluted samples were mixed with a constant amount of SARS-CoV-2 and incubated for 1 h at 37 °C. The antibody-virus-mix was then directly applied to each well (n = 3 per dilution) and incubated for 24 h at 37 °C. The infectious dose for each virus was pre-determined on VeroE6_UNC_ cells to yield 50-60% antigen-positive cells upon this incubation period (USA-WA1/2020: 1,250 PFU/well and B.1.351: 175 PFU/well). Cells were subsequently fixed by adding an equal volume of 7% formaldehyde to the wells, followed by permeabilization with 0.1% Triton X-100 for 10 min. After extensive washing, cells were incubated for 1h at RT with blocking solution of 5% goat serum in PBS (Jackson ImmunoResearch, catalog no. 005–000-121). A rabbit polyclonal anti-SARS-CoV-2 nucleocapsid antibody (GeneTex, catalog no. GTX135357) was added to the cells at 1:1,000 dilution in blocking solution and incubated overnight at 4 °C. Next, goat anti-rabbit AlexaFluor 594 (Life Technologies, catalog no. A-11012) was used as a secondary antibody at a dilution of 1:2,000 and incubated overnight at 4 °C. Nuclei were stained with Hoechst 33342 (ThermoFisher Scientific, catalog no. 62249) at a 1 µg/mL. Images were acquired with a fluorescence microscope and analyzed using ImageXpress Micro XLS (Molecular Devices, Sunnyvale, CA). All statistical analyses were done using Prism 8 software (Graphpad).

### Biotinylation of viral protein for use in flow cytometry

Purified and Avi-tagged SARS-CoV-2 NTD was biotinylated using the Biotin-Protein Ligase-BIRA kit according to manufacturer’s instructions (Avidity) as described before (Robbiani et al., 2020). Ovalbumin (Sigma, A5503-1G) was biotinylated using the EZ-Link Sulfo-NHS-LC-Biotinylation kit according to the manufacturer’s instructions (Thermo Scientific). Biotinylated ovalbumin was conjugated to streptavidin-BV711 (BD biosciences, 563262), Gamma NTD to streptavidin-PE (BD Biosciences, 554061) and Wuhan-Hu-1 NTD to streptavidin-AF647 (Biolegend, 405237).

### Flow cytometry and single cell sorting

Single cell sorting by flow cytometry was performed as described (Robbiani et al., 2020). Briefly, peripheral blood mononuclear cells were enriched for B cells by negative selection using a pan-B-cell isolation kit according to the manufacturer’s instructions (Miltenyi Biotec, 130-101-638). The enriched B cells were incubated in FACS buffer (1× PBS, 2% FCS, 1 mM EDTA) with the following anti-human antibodies (all at 1:200 dilution): anti-CD20-PECy7 (BD Biosciences, 335793), anti-CD3-APC-eFluro 780 (Invitrogen, 47-0037-41), anti-CD8-APC-eFluor 780 (Invitrogen, 47-0086-42), anti-CD16-APC-eFluor 780 (Invitrogen, 47-0168-41), anti-CD14-APC-eFluor 780 (Invitrogen, 47-0149-42), as well as Zombie NIR (BioLegend, 423105) and fluorophore-labelled RBD and ovalbumin (Ova) for 30 min on ice. Single CD3−CD8−CD14−CD16−CD20+Ova−Gamma NTD-PE^+^Wuhan-Hu-1 NTD-AF647^+^ B cells were sorted into individual wells of 96-well plates containing 4 μl of lysis buffer (0.5× PBS, 10 mM DTT, 3,000 units/ml RNasin Ribonuclease Inhibitors (Promega, N2615) per well using a FACS Aria III and FACSDiva software (Becton Dickinson) for acquisition and FlowJo for analysis. The sorted cells were frozen on dry ice, and then stored at −80 °C or immediately used for subsequent RNA reverse transcription.

### Antibody sequencing, cloning and expression

Antibodies were identified and sequenced as described previously (Robbiani et al., 2020). In brief, RNA from single cells was reverse-transcribed (SuperScript III Reverse Transcriptase, Invitrogen, 18080-044) and the cDNA stored at −20 °C or used for subsequent amplification of the variable IGH, IGL and IGK genes by nested PCR and Sanger sequencing. Sequence analysis was performed using MacVector. Amplicons from the first PCR reaction were used as templates for sequence- and ligation-independent cloning into antibody expression vectors. Recombinant monoclonal antibodies and Fabs were produced and purified as previously described (Robbiani et al., 2020).

### Biolayer interferometry

BLI assays were performed on the Octet Red instrument (ForteBio) at 30 °C with shaking at 1,000 r.p.m. Epitope binding assays were performed with protein A biosensor (ForteBio 18-5010), following the manufacturer’s protocol “classical sandwich assay” as follows: (1) Sensor check: sensors immersed 30 sec in buffer alone (buffer ForteBio 18-1105), (2) Capture 1st Ab: sensors immersed 10 min with Ab1 at 10 µg/mL, (3) Baseline: sensors immersed 30 sec in buffer alone, (4) Blocking: sensors immersed 5 min with IgG isotype control at 10 µg/mL. (5) Baseline: sensors immersed 30 sec in buffer alone, (6) Antigen association: sensors immersed 5 min with NTD at 10 µg/mL. (7) Baseline: sensors immersed 30 sec in buffer alone. (8) Association Ab2: sensors immersed 5 min with Ab2 at 10 µg/mL. Curve fitting was performed using the Fortebio Octet Data analysis software (ForteBio). Affinity measurement of anti-SARS-CoV-2 IgGs binding were corrected by subtracting the signal obtained from traces performed with IgGs in the absence of WT NTD. The kinetic analysis using protein A biosensor (ForteBio 18-5010) was performed as follows: (1) baseline: 60sec immersion in buffer. (2) loading: 200sec immersion in a solution with IgGs 10 μg/ml. (3) baseline: 200sec immersion in buffer. (4) Association: 300sec immersion in solution with WT NTD at 200, 100, 50 or 25 μg/ml (5) dissociation: 600sec immersion in buffer. Curve fitting was performed using a fast 1:1 binding model and the Data analysis software (ForteBio). Mean *K*_D_ values were determined by averaging all binding curves that matched the theoretical fit with an R^2^ value ≥ 0.8.

### Recombinant protein expression for structural studies

Expression and purification of stabilized SARS-CoV-2 6P ectodomain was conducted as previously described(Barnes et al., 2020a). Briefly, constructs encoding the SARS-CoV-2 S ectodomain (residues 16-1206 with 6P stabilizing mutations(Hsieh et al., 2020), a mutated furin cleavage site, and C-terminal foldon trimerization motif followed by hexa-His tag) were used to transiently transfect Expi293F cells (Gibco). Four days after transfection, supernatants were harvested and S 6P proteins were purified by nickel affinity and size-exclusion chromatography. Peak fractions from size-exclusion chromatography were identified by SDS-PAGE, and fractions corresponding to spike trimers were pooled and stored at 4°C. Fabs and IgGs were expressed, purified, and stored as previously described(Barnes et al., 2020a).

### Cryo-EM sample preparation

Purified Fabs were mixed with SARS-CoV-2 S 6P trimer at a 1.1:1 molar ratio of Fab-to-protomer for 30 minutes at room temperature. Fab-S complexes were concentrated to 3-4 mg/mL prior to deposition on a freshly glow-discharged 300 mesh, 1.2/1.3 Quantifoil grid (Electron Microscopy Sciences). Immediately prior to deposition of 3 µL of complex onto grid, fluorinated octyl-maltoside (Anatrace) was added to the sample to a final concentration of 0.02% w/v. Samples were vitrified in 100% liquid ethane using a Mark IV Vitrobot (Thermo Fisher) after blotting at 22°C and 100% humidity for 3s with Whatman No. 1 filter paper.

### Cryo-EM data collection and processing

Single-particle cryo-EM data were collected on a Titan Krios transmission electron microscope (Thermo Fisher) equipped with a Gatan K3 direct detector, operating at 300 kV and controlled using SerialEM automated data collection software(Mastronarde, 2005). A total dose of ∼60 e^-^ /Å^2^ was accumulated on each movie comprising 40 frames with a pixel size of 0.515 Å (C1520-S dataset) or 0.852 Å (C1717-S and C1791-S) and a defocus range of −1.0 and −2.5 µm. Further data collection parameters are summarized in Table S5.

Movie frame alignment, CTF estimation, particle-picking and extraction were carried out using cryoSPARC v3.1(Punjani et al., 2017). Reference-free particle picking and extraction were performed on dose-weighted micrographs curated to remove images with poor CTF fits or signs of crystalline ice. A subset of 4x-downsampled particles were used to generate *ab initio* models, which were then used for heterogeneous refinement of the entire dataset in cryoSPARC. Particles belonging to classes that resembled Fab-S structures were extracted, downsampled x2 and subjected to 2D classification to select well-defined particle images. 3D classifications (k=6, tau_fudge=4) were carried out using Relion v3.1.1(Fernandez-Leiro and Scheres, 2017) without imposing symmetry and a soft mask. Particles corresponding to selected classes were re-extracted without binning and 3D refinements were carried out using non-uniform refinement in cryoSPARC. Particle stacks were split into individual exposure groups based on the beamtilt angle used for data collection and subjected to per particle CTF refinement and aberration corrections. Another round of non-uniform refinement in cryoSPARC was then performed. For focused classification and local refinements of the Fab V_H_V_L_-NTD interface, particles were 3D classified in Relion without alignment using a mask that encompassed the Fab-NTD region. Particles in good 3D classes were then used for local refinement in cryoSPARC. Details of overall resolution and locally-refined resolutions according to the gold-standard Fourier shell correlation of 0.143 criterion(Bell et al., 2016) can be found in Table S5.

### Cryo-EM structure modeling, refinement, and analyses

Coordinates for initial complexes were generated by docking individual chains from reference structures (see Table S5) into cryo-EM density using UCSF Chimera(Goddard et al., 2007). Initial models for Fabs were generated from coordinates from PDB 6RCO (for C1717 Fab) or PDB 7RKS (for C5120). Models were refined using one round of rigid body refinement followed by real space refinement in Phenix(Adams et al., 2010). Sequence-updated models were built manually in Coot(Emsley et al., 2010) and then refined using iterative rounds of refinement in Coot and Phenix. Glycans were modeled at potential *N*-linked glycosylation sites (PNGSs) in Coot. Validation of model coordinates was performed using MolProbity(Chen et al., 2010).

Structure figures were made with UCSF ChimeraX(Goddard et al., 2018). Local resolution maps were calculated using cryoSPARC v3.1(Punjani et al., 2017). Buried surface areas were calculated using PDBePISA(Krissinel and Henrick, 2007) and a 1.4Å probe. Potential hydrogen bonds were assigned as interactions that were <4.0Å and with A-D-H angle >90°. Potential van der Waals interactions between atoms were assigned as interactions that were <4.0Å. Hydrogen bond and van der Waals interaction assignments are tentative due to resolution limitations. Spike epitope residues were defined as residues containing atom(s) within 4Å of a Fab atom for the C1520-S and C1717-S complexes, and defined as spike C*α* atom within 7Å of a Fab C*α* atom for the C1791-S complex.

### Computational analyses of antibody sequences

Antibody sequences were trimmed based on quality and annotated using Igblastn v.1.14. with IMGT domain delineation system. Annotation was performed systematically using Change-O toolkit v.0.4.540 (Gupta et al., 2015). Heavy and light chains derived from the same cell were paired, and clonotypes were assigned based on their V and J genes using in-house R and Perl scripts (Figure S4). All scripts and the data used to process antibody sequences are publicly available on GitHub (https://github.com/stratust/igpipeline/tree/igpipeline2_timepoint_v2).

The frequency distributions of human V genes in anti-SARS-CoV-2 antibodies from this study was compared to 131,284,220 IgH and IgL sequences generated by (Soto et al., 2019) and downloaded from cAb-Rep(Guo et al., 2019), a database of human shared BCR clonotypes available at https://cab-rep.c2b2.columbia.edu/. Based on the 174 distinct V genes that make up the 3085 analyzed sequences from Ig repertoire of the 6 participants present in this study, we selected the IgH and IgL sequences from the database that are partially coded by the same V genes and counted them according to the constant region. The frequencies shown in (Figures 2D and S5) are relative to the source and isotype analyzed. We used the two-sided binomial test to check whether the number of sequences belonging to a specific IgHV or IgLV gene in the repertoire is different according to the frequency of the same IgV gene in the database. Adjusted p-values were calculated using the false discovery rate (FDR) correction. Significant differences are denoted with stars.

Nucleotide somatic hypermutation and CDR3 length were determined using in-house R and Perl scripts. For somatic hypermutations, IGHV and IGLV nucleotide sequences were aligned against their closest germlines using Igblastn and the number of differences were considered nucleotide mutations. The average mutations for V genes were calculated by dividing the sum of all nucleotide mutations across all participants by the number of sequences used for the analysis. To calculate the GRAVY scores of hydrophobicity (Kyte and Doolittle, 1982) we used Guy H.R. Hydrophobicity scale based on free energy of transfer (kcal/mole) (Guy, 1985) implemented by the R package Peptides (the Comprehensive R Archive Network repository; https://journal.r-project.org/archive/2015/RJ-2015-001/RJ-2015-001.pdf). We used heavy chain CDR3 amino acid sequences from this study and 22,654,256 IGH CDR3 sequences from the public database of memory B cell receptor sequences (DeWitt et al., 2016). The two-tailed Wilcoxon nonparametric test was used to test whether there is a difference in hydrophobicity distribution.

## SUPPLEMENTARY FIGURES AND TABLES

**Figure S1.**
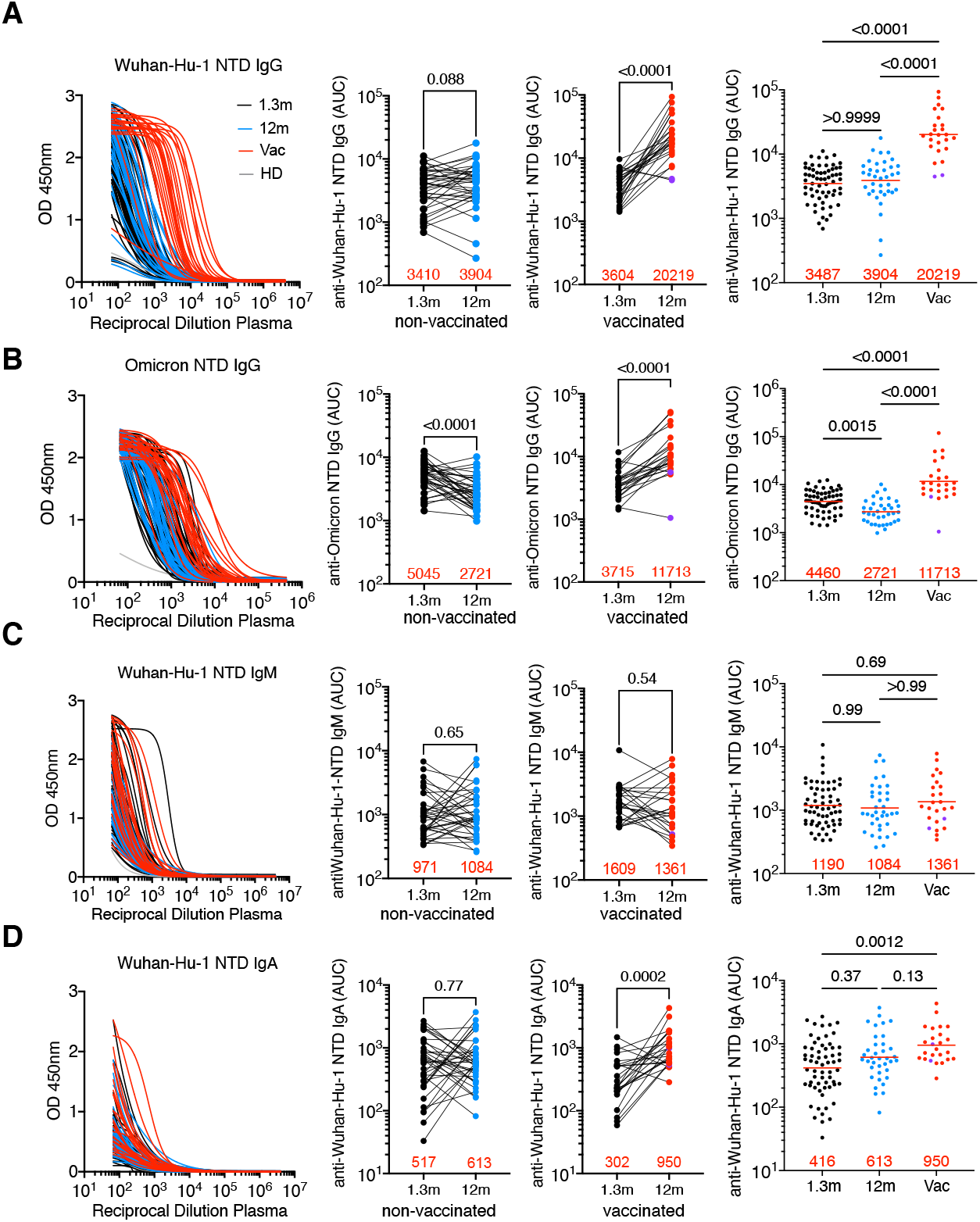
Plasma anti-SARS-CoV-2 NTD reactivity. **A-B**, Plasma IgG antibody binding to Wuhan-Hu-1 NTD (**A**) and Omicron NTD protein (**B**) in unvaccinated and vaccinated (vac) convalescent individuals at 1.3 month(Robbiani et al., 2020) and 12 months(Wang et al., 2021c) after infection (n=62). Graphs showing ELISA curves from individuals 1.3 months after infection (black lines), from individuals 12m after infection unvaccinated (blue lines) and vaccinated (red lines) (left panels). Area under the curve (AUC) over time in non-vaccinated and vaccinated individuals, as indicated (middle panels). Lines connect longitudinal samples. Two outliers who received their first dose of vaccine 24-48 hours before sample collection are depicted in purple. Numbers in red indicate geometric mean AUC at the indicated timepoint. **C-D**, same as **A**, shown as IgM (**C**) and IgA (**D**) antibody binding to SARS-CoV-2 NTD 1.3- and 12-months after infection. Statistical significance was determined using Wilcoxon matched-pairs signed rank tests. Right panel shows combined values as a dot plot for all individuals. Statistical significance was determined through the Kruskal Wallis test with subsequent Dunn’s multiple comparisons.

**Figure S2.**
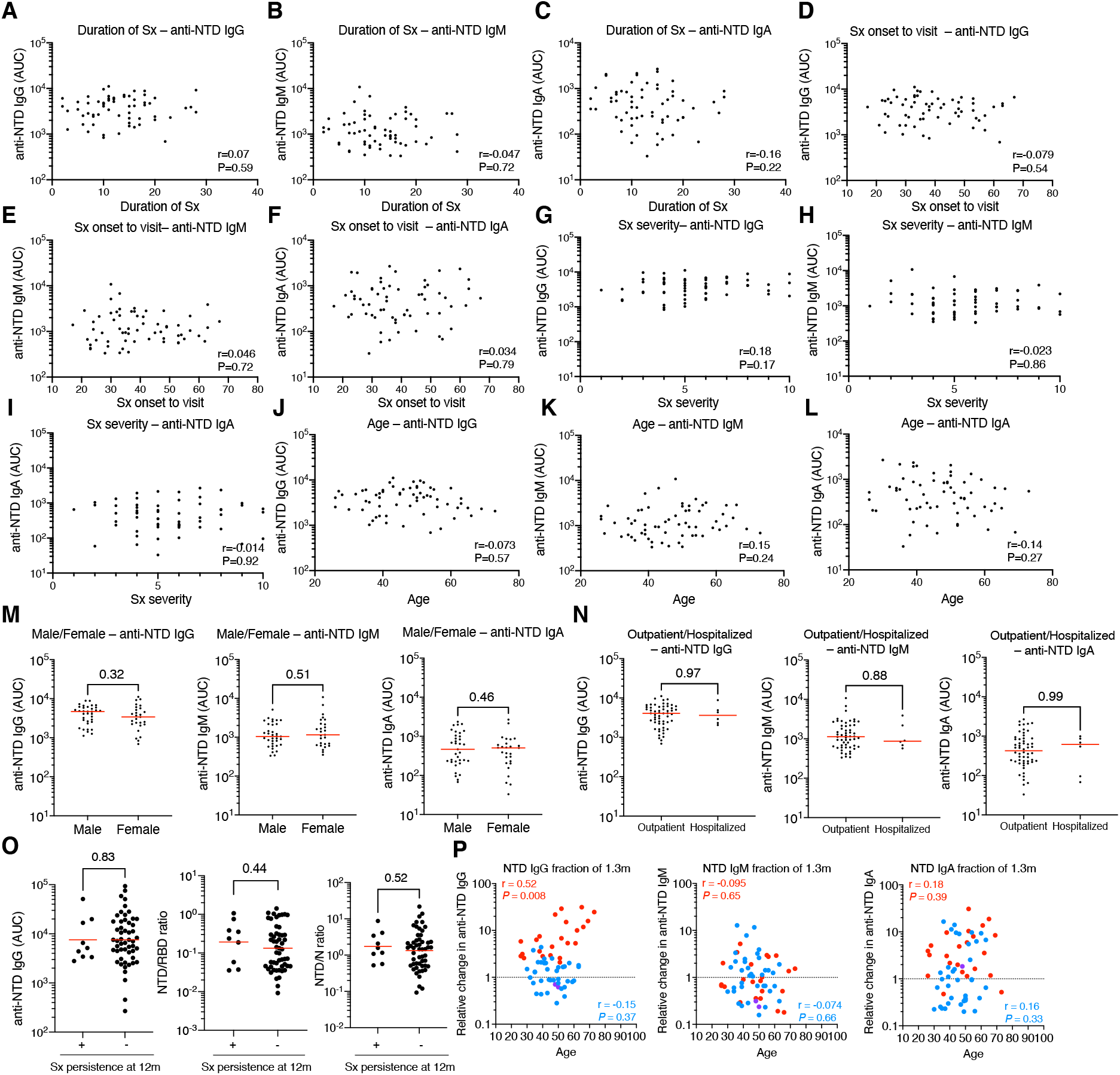
Clinical correlations of plasma anti-SARS-CoV-2 NTD antibody titers. **A-C**, Clinical correlations of plasma anti-NTD IgG titers at 1.3month after infection(Robbiani et al., 2020). **A-C**, Duration of Symptom (Sx) in days (X axis) plotted against normalized AUC for plasma IgG (**A**), IgM (**B**) and IgA (**C**) binding to NTD (Y axis). **D-F**, Sx onset to time of sample collection in days plotted against normalized AUC for plasma IgG (**D**), IgM (**E**) and IgA (**F**) anti-NTD. **G-I**, Sx severity plotted against normalized AUC for plasma IgG (**G**), IgM (**H**) and IgA (**I**) anti-NTD. **J-L**, Age plotted against normalized AUC for plasma IgG (**J**), IgM (**K**) and IgA(**L**) anti-NTD. **M.** Normalized plasma IgG, IgM and IgA AUC for males (n=34) and females (n=28). **N**, Normalized plasma IgG, IgM and IgA AUC for outpatient (n = 56) and hospitalized (n = 6). **O**, Anti-NTD IgG, NTD/RBD IgG ratio and NTD/N IgG ratio at 12 months post-infection in individuals reporting persistent symptoms (+) compared to individuals who are symptom-free (-) 12 months post-infection. **P**, Correlation of remaining plasma titers at 12 months (expressed as the fraction of 1.3-month titers on the Y axis) and participant age for anti-NTD IgG (left), anti-NTD IgM (middle), anti-NTD IgA (right). For **A-L, P**, the correlations were analyzed by two-tailed Spearman’s tests; For **M-N**, Statistical significance was determined using two-tailed Mann– Whitney U-tests.

**Figure S3.**
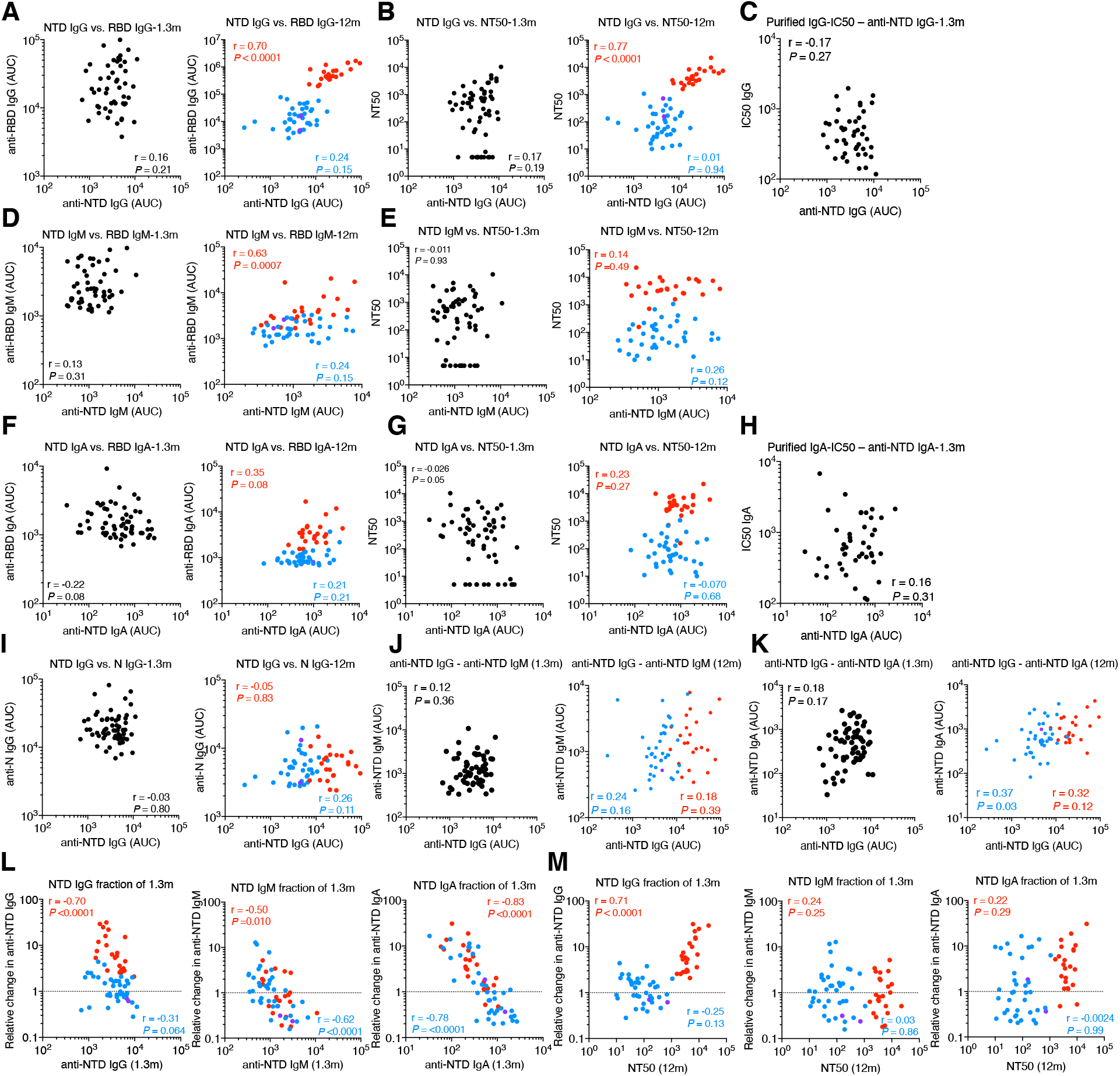
Plasma activity. **A,** Correlation of 1.3- (left) and 12-month (right) titers of anti-NTD IgG and anti-RBD IgG at each timepoint, separately (Robbiani et al., 2020; Wang et al., 2021c). **B**, Correlation of 1.3-(left) and 12-month (right) titers of anti- NTD IgG and NT_50_ at each timepoint, separately (Robbiani et al., 2020; Wang et al., 2021c). **C**, Correlation of 1.3-month plasma titers of anti-NTD IgG and purified plasma IgG IC_50_ (Wang et al., 2021b). **D**, Correlation of 1.3-(left) and 12-month (right) plasma titers of anti-NTD IgM and anti-RBD IgM (Robbiani et al., 2020; Wang et al., 2021c). **E**, Correlation of 1.3-(left) and 12-month (right) plasma titers of anti-NTD IgM and NT_50_ (Robbiani et al., 2020; Wang et al., 2021c). **F**, Correlation of 1.3-(left) and 12-month (right) plasma titers of anti-NTD IgA and anti-RBD IgA(Robbiani et al., 2020; Wang et al., 2021c). **G**, Correlation of 1.3- (left) and 12-month (right) plasma titers of anti-NTD IgA and NT_50_ (Robbiani et al., 2020; Wang et al., 2021c). **H**, Correlation of 1.3-month plasma titers of anti-NTD IgA and purified plasma IgA IC_50_ (Wang et al., 2021b). **I**, Correlation of 1.3- (left) and 12-month (right) plasma titers of anti-NTD IgG and anti-N IgG (Robbiani et al., 2020; Wang et al., 2021c). **J**, Correlation of 1.3- (left) and 12-month (right) plasma titers of anti-NTD IgG and anti-NTD IgM. **K**, Correlation of 1.3- (left) and 12-month (right) plasma titers of anti-NTD IgG and anti-NTD IgA. **L**, Correlation of remaining plasma titers at 12 months (expressed as the fraction of 1.3-month titers on the Y axis) and 1.3- month plasma titers of anti-NTD IgG (left), anti-NTD IgM (middle), anti-NTD IgA (right) (Robbiani et al., 2020; Wang et al., 2021c). **M**, Correlation of remaining plasma titers of anti-NTD IgG (left), anti-NTD IgM (middle), anti-NTD IgA (right) (expressed as the fraction of 1.3-month titers on the Y axis) and plasma NT_50_ at 12 months (Wang et al., 2021c). For **A**, **B**, **D**, **E**, **F**, **G**, **I-M**, graphs showing dot plots from individuals 1.3 months after infection (black), non-vaccinated convalescents (blue) or vaccinated convalescents (red) 12m after infection. Statistical significance was determined using the Spearman correlation test for the non-vaccinated and vaccinated subgroups independently. All experiments were performed at least in duplicate.

**Figure S4.**
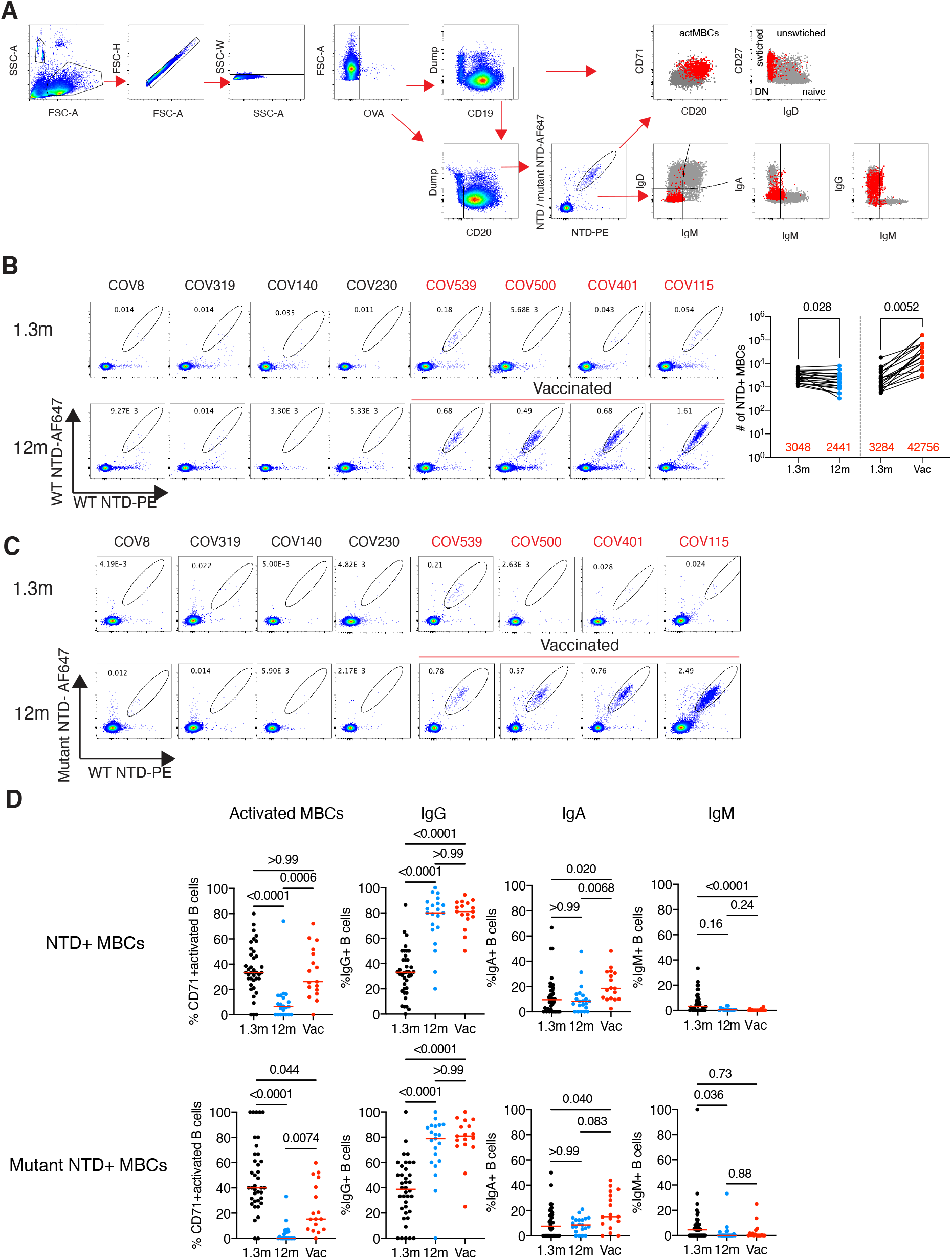
Flow cytometry. **A**, Gating strategy. Gating was on singlets that were CD20^+^ or CD19^+^ and CD3^-^CD8^-^CD16^-^Ova^-^. Anti-IgG, IgM, IgA, IgD, CD71 and CD27 antibodies were used for B cell phenotype analysis. **B** and **C**, Flow cytometry showing the percentage of NTD-double positive (**B**) and AF647-mutant NTD cross-reactive (**C**) memory B cells from 1.3- and 12-months post-infection in 39 selected patients. **D**. B cell phenotyping. Expression of CD71, IgG, IgA, IgM in antigen-specific memory B cells. Each dot is one individual. Red horizontal bars indicate mean values. Statistical significance was determined using two-tailed Mann–Whitney U-tests (**B**) and Kruskal Wallis test with subsequent Dunn’s multiple comparisons (**D**).

**Figure S5.**
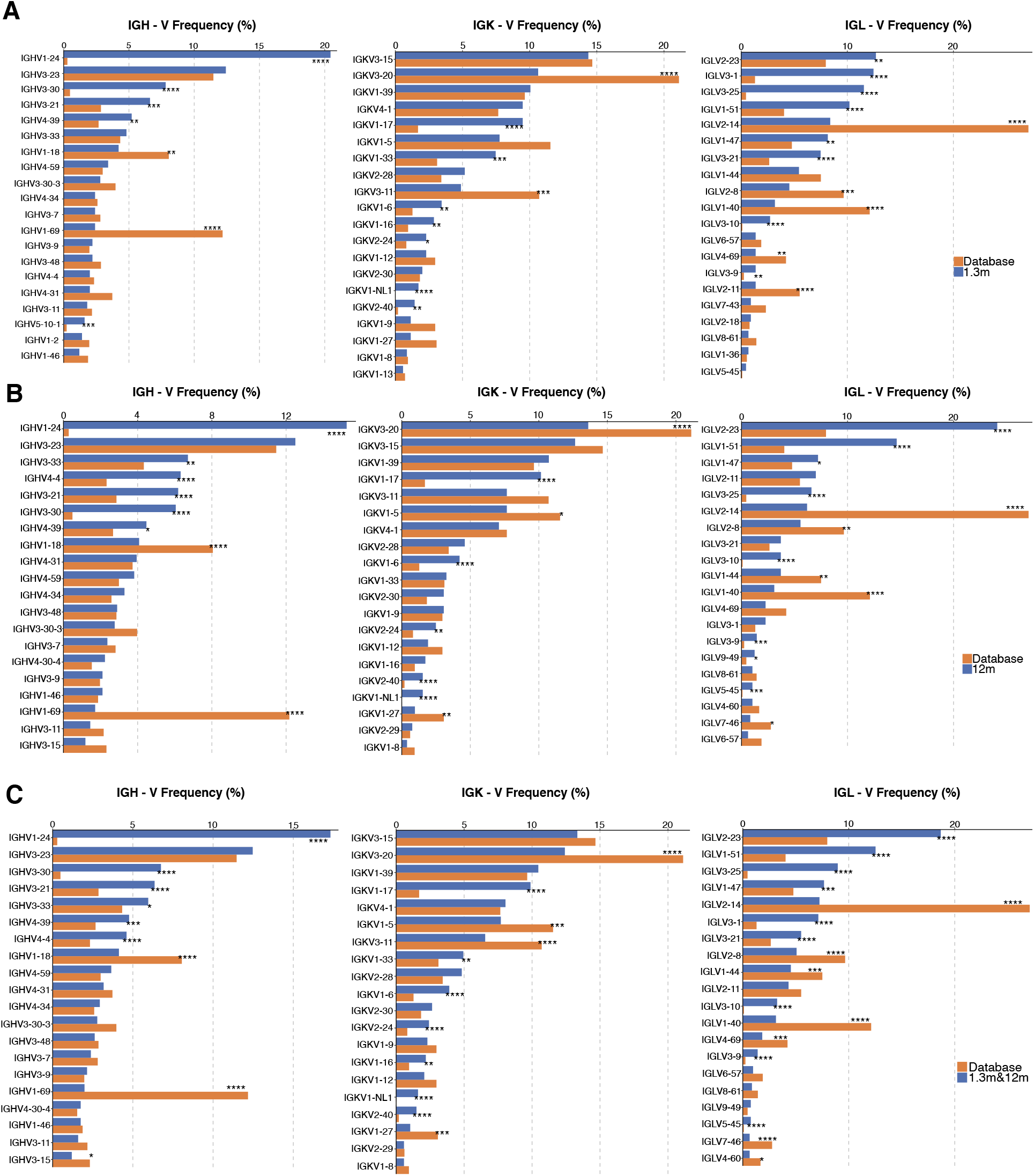
Frequency distribution of human V genes. Comparison of the frequency distribution of human V genes for heavy chain and light chains of anti-NTD antibodies from this study and from a database of shared clonotypes of human B cell receptor generated by Cinque Soto et al(Soto et al., 2019). **A** and **B**, Graph shows relative abundance of human IGVH, IGVK and IGVL genes Sequence Read Archive accession SRP010970 (orange), 1.3- (**A**), and 12-month antibodies (**B**). **C**, Same as **A** and **B** but shows the comparison between Sequence Read Archive accession SRP010970 (orange) and combined (1.3m and 12m) anti-NTD antibodies (blue). Statistical significance was determined by two-sided binomial test. Significant differences are denoted with stars.

**Figure S6.**
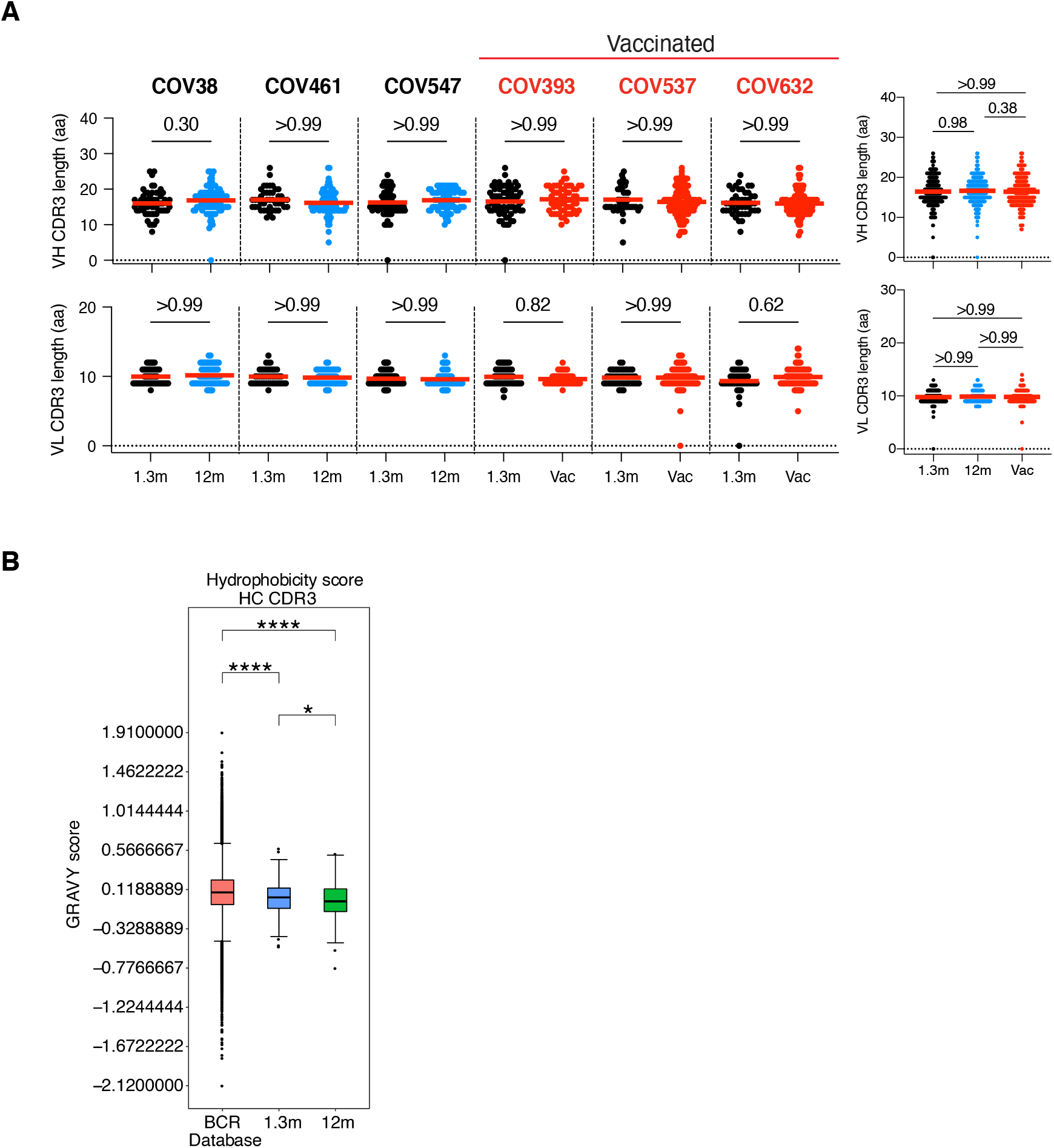
CDR3 length and hydrophobicity of anti-SARS-CoV-2 NTD antibodies. **A**, IGVH and IGVL CDR3s length (Y axis) for anti-SARS-CoV-2 NTD antibodies isolated from 6 individuals 1.3- and 12-months after infection. Vaccinees are marked in red. Each dot represents one antibody from individuals 1.3 months after infection (black), non-vaccinated convalescents (blue) or vaccinated convalescents (red) 12m after infection. Statistical significance was determined using Kruskal Wallis test with subsequent Dunn’s multiple comparisons. **B**, Distribution of the hydrophobicity GRAVY scores at the IGH CDR3 in antibody sequences from this study compared to a public database (see Methods for statistical analysis). The box limits are at the lower and upper quartiles, the center line indicates the median, the whiskers are 1.5x interquartile range and the dots represent outliers.

**Figure S7.**
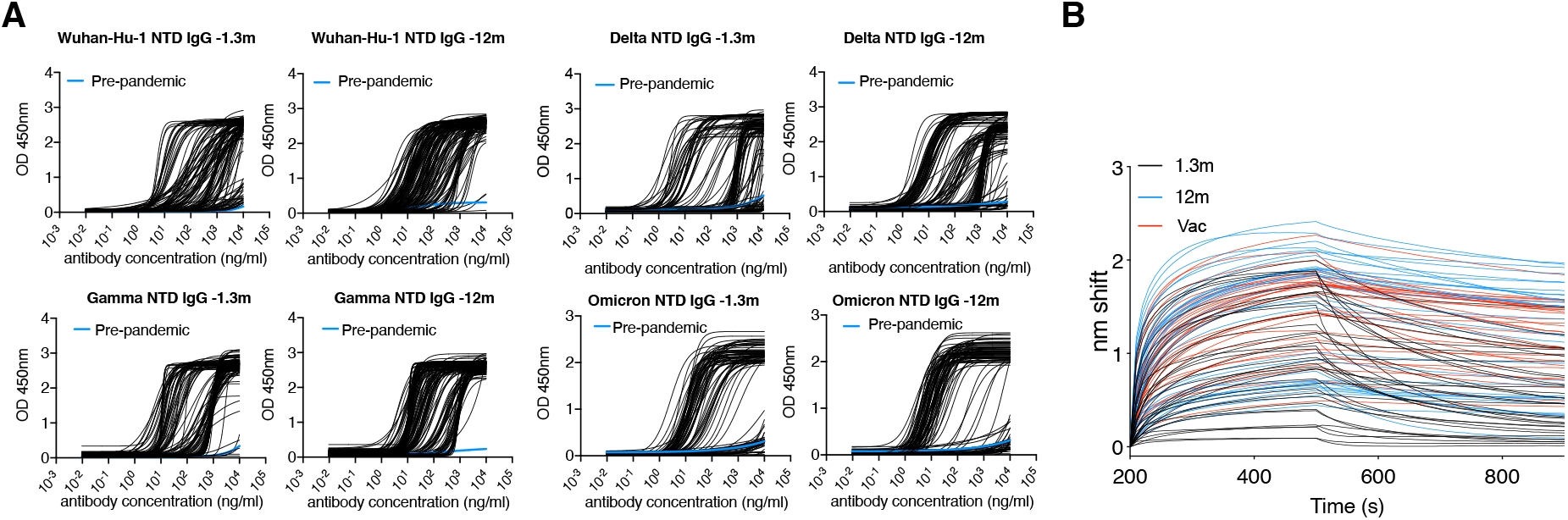
Binding curves of anti-SARS-CoV-2 NTD monoclonal antibodies. **A**, ELISA binding curves of mAbs isolated from convalescents individuals at 1.3- and 12 months after infection, EC_50_s against SARS-CoV-2 Wuhan-Hu-1-, Delta-, Gama- or Omicron-NTD were shown. **B**, Biolayer interferometry association and dissociation curves for 1.3- (black), 12-month convalescents (blue lines), and 12-month convalescent vaccinees (red lines).

**Figure S8.**
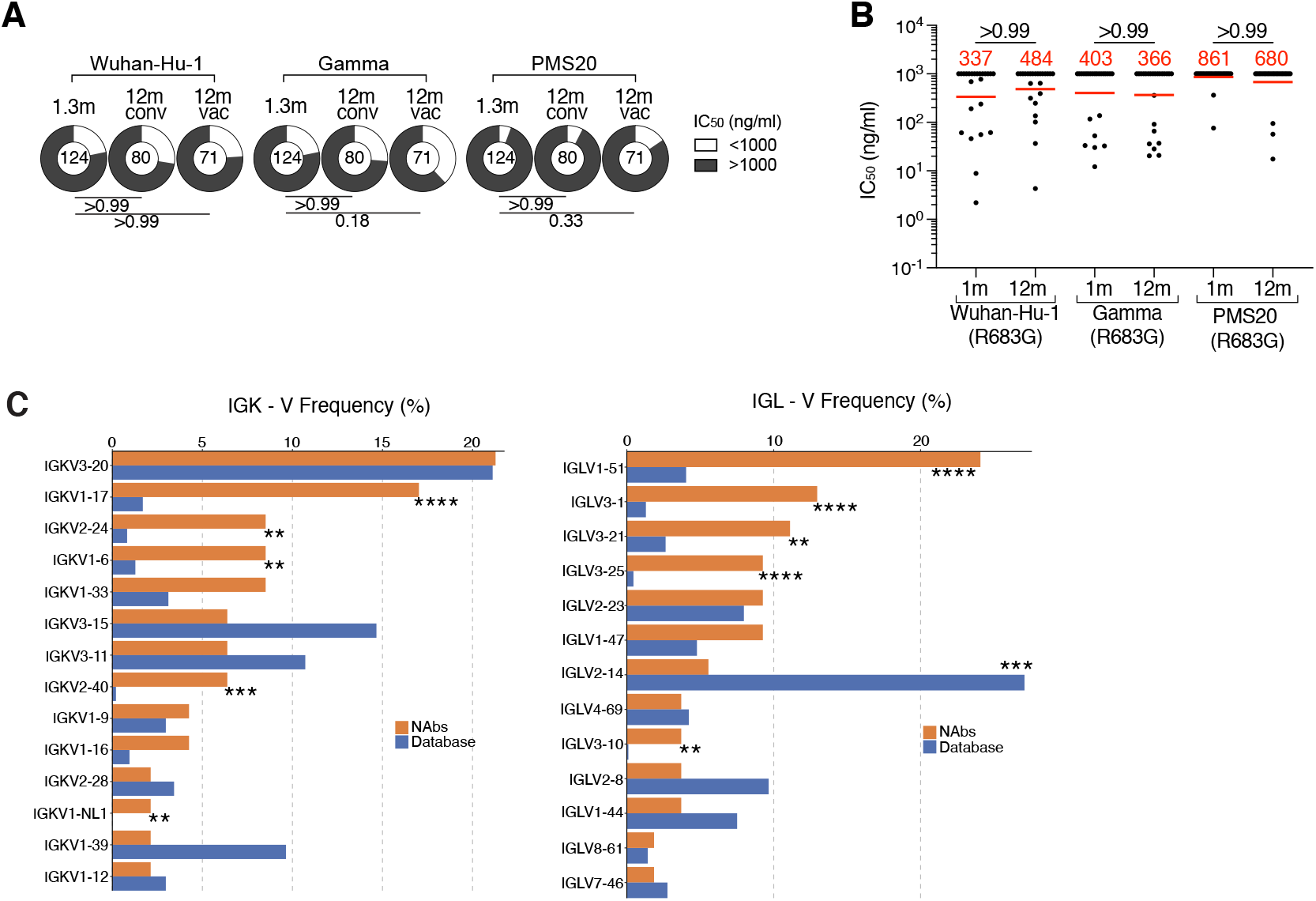
anti-SARS-CoV-2 NTD monoclonal antibodies. **A**, Corresponding to Figure 3A, Pie charts illustrate the fraction of neutralizing (white slices) and non-neutralizing (dark grey slices) antibodies, inner circle shows the number of antibodies tested per group. Statistical significance was determined with Fisher’s exact test with subsequent Bonferroni-Dunn correction. **B**, Graph shows anti-SARS-CoV-2 neutralizing activity of shared clones of antibodies isolated 1.3- and 12-months after infection, measured by a SARS-CoV-2 pseudovirus neutralization assay(Robbiani et al., 2020; Schmidt et al., 2020). Statistical significance was determined through Friedman test with subsequent Dunn’s multiple comparisons. Horizontal bars and red numbers indicate geometric mean values. **C**, Graph shows comparison of the frequency distributions of human IGK and IGL V genes of anti-SARS-CoV-2 NTD neutralizing antibodies from donors at 1.3 month(Robbiani et al., 2020) and 12 months(Wang et al., 2021c) after infection. Statistical significance was determined by two-sided binomial test.

**Figure S9.**
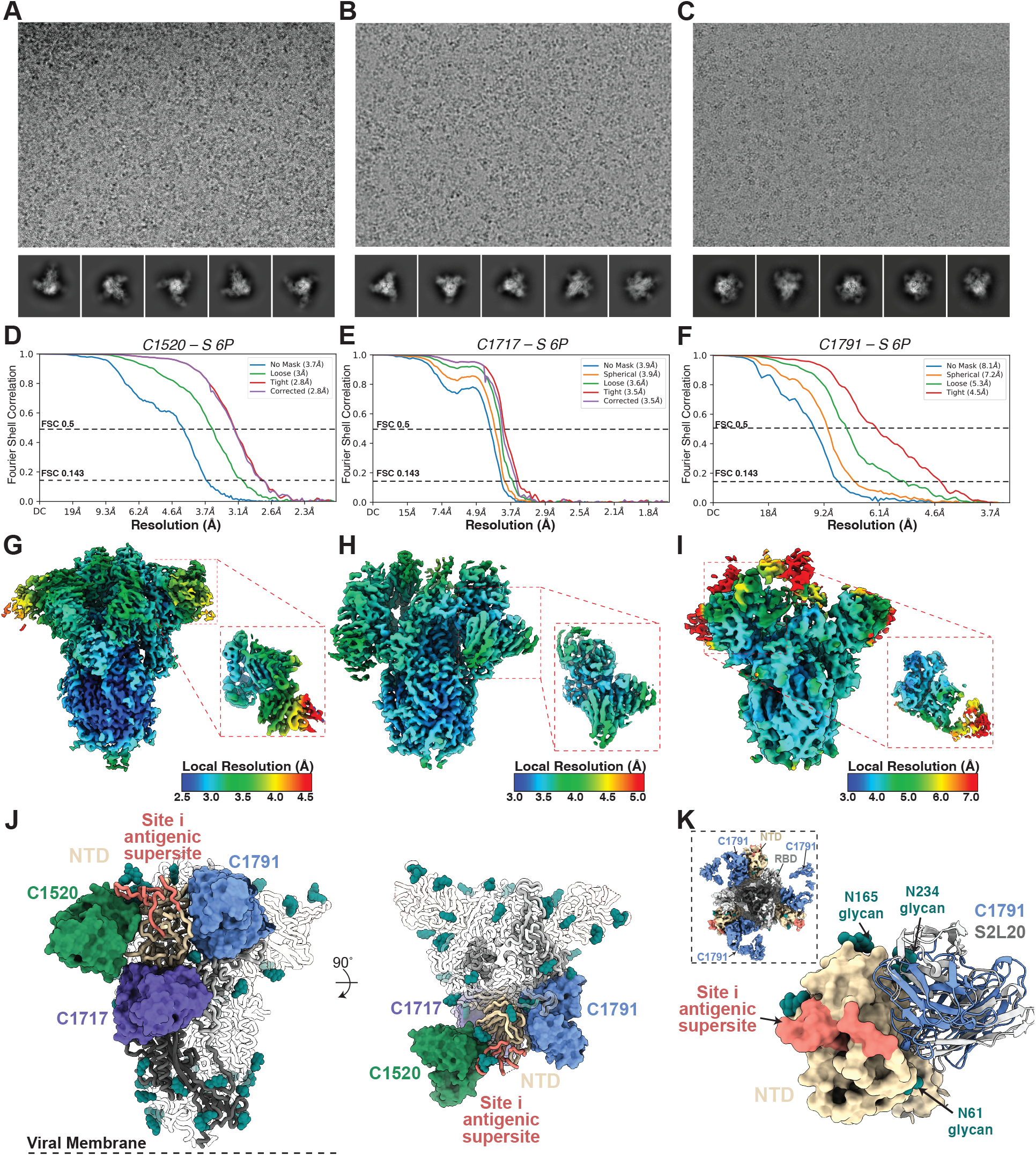
Cryo-EM data processing and validation. **A-F,** Representative micrographs selected from total datasets (see Table S5), 2D class averages, and gold-standard FSC plots for **A,D** C1520-S 6P; **B,E** C1717-S 6P; and **C,F** C1791-S 6P. **G-I,** Local resolution estimations for **G,** C1520-S, **H,** C1717-S, and **I,** C1791-S complexes. Insets show resolution estimates for Fab-NTD focused refinements. **J,** Composite model illustrating non-overlapping epitopes of NTD-targeting mAbs (C1520 – green; C1717 – purple; C1791 – slate blue) bound to a single NTD (wheat) on a spike trimer. The site i antigenic supersite (coral) is shown as reference. **K,** Structural superposition of NTD-targeting antibodies C1791 (slate blue) and S2L20 (gray – PDB 7N8I) on a NTD (wheat surface rendering). The NTD site i antigenic supersite is depicted as in panel **J**. Cryo-EM density for the C1791-S is also shown (inset).

**Figure S10.**
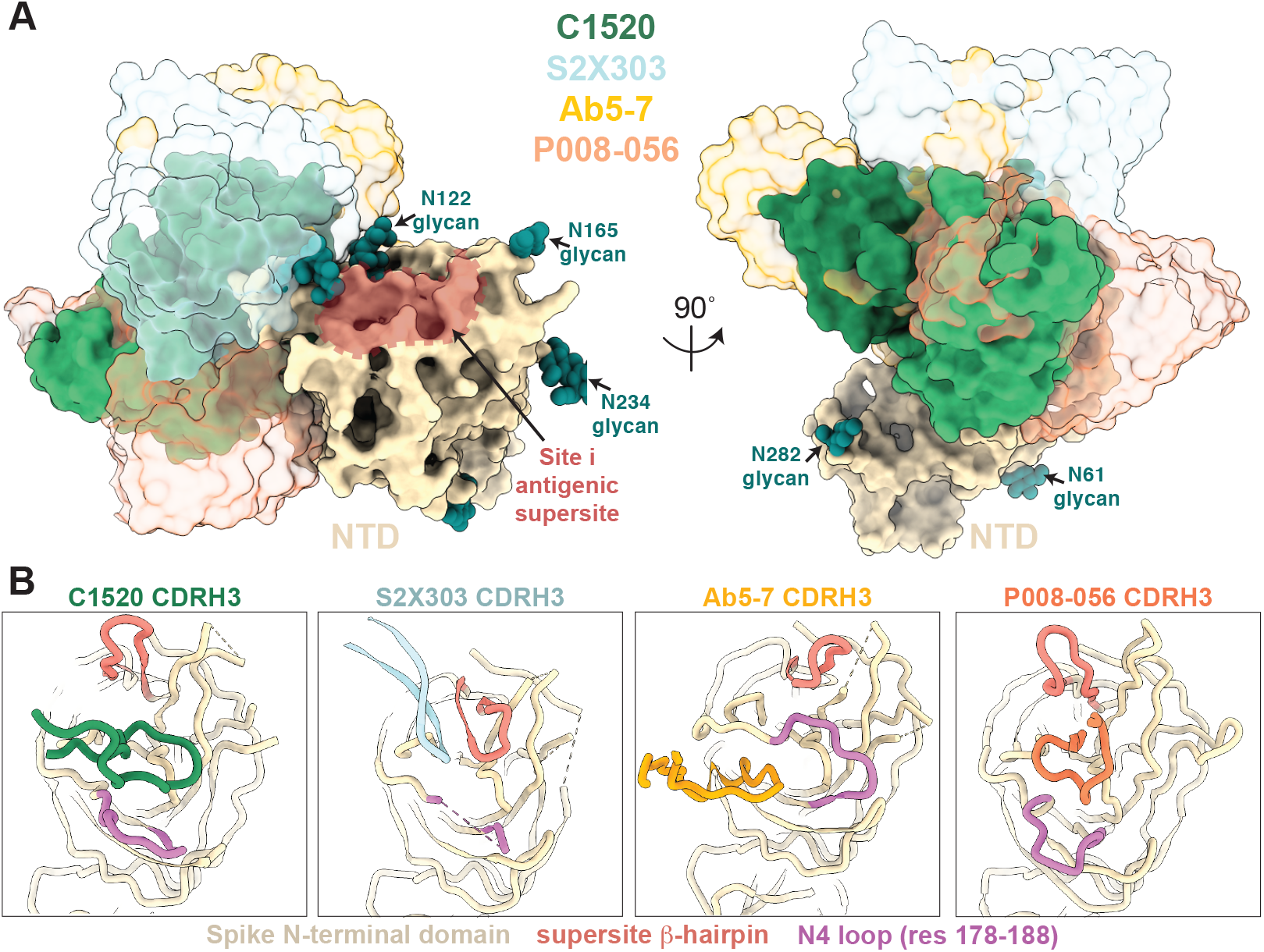
Comparison of C1520-like NTD neutralizing antibodies. **A,** Overlay of V_H_ and V_L_ domains of C1520 (green), S2X303 (PDB 7SOF; powder blue), Ab5-7 (PDB 7RW2; gold), and P008-056 (PDB 7NTC; orange) after alignment on NTD residues 27-303, illustrating distinct binding poses to NTD epitopes adjacent to the site i supersite (coral). **B,** Comparison of CDRH3 loop position, the NTD supersite b-hairpin (coral), and NTD N4-loop (orchid) for antibodies shown in panel **A**.

## SUPPLEMENTARY TABLES

**<u>Table S1</u>: Individual participant characteristics**

**<u>Table S2</u>: Antibody sequences from patients is provided as a separate Excel file.**

**<u>Table S3</u>: Sequences, half maximal effective concentrations (EC50s) and inhibitory concentrations (IC50s) of the cloned monoclonal antibodies is provided as a separate Excel file.**

**<u>Table S4</u>: V gene usage and neutralization activity of anti-NTD nAbs.**

**<u>Table S5</u>: Cryo-EM data collection and processing statistics**

